# Sperm migration in the genital tract - *in silico* experiments identify key factors for reproductive success

**DOI:** 10.1101/2020.03.05.977967

**Authors:** Jorin Diemer, Jens Hahn, Björn Goldenbogen, Karin Müller, Edda Klipp

## Abstract

Sperm migration in the female genital tract controls sperm selection and, therefore, reproductive success as male gametes are conditioned for fertilization while their number is dramatically reduced. Mechanisms underlying sperm migration are mostly unknown, since *in vivo* investigations are mostly unfeasible for ethical or practical reasons. By presenting a spatio-temporal model of the mammalian female genital tract combined with agent-based description of sperm motion and interaction as well as parameterizing it with bovine data, we offer an alternative possibility for studying sperm migration *in silico*. The model incorporates genital tract geometry as well as biophysical principles of sperm motion observed *in vitro* such as positive rheotaxis and thigmotaxis. This model for sperm migration from vagina to oviducts was successfully tested against *in vivo* data from literature. We found that physical sperm characteristics such as velocity and directional stability as well as sperm-fluid interactions and wall alignment are critical for success, i.e. sperms reaching the oviducts. Therefore, we propose that these identified sperm parameters should be considered in detail for conditioning sperm in artificial selection procedures since the natural processes are normally bypassed in reproductive *in vitro* technologies. The tremendous impact of mucus flow to support sperm accumulation in the oviduct highlights the importance of a species-specific optimum time window for artificial insemination regarding ovulation. Predictions from our extendable *in silico* experimental system will improve assisted reproduction in humans, endangered species, and livestock.

## Introduction

Mammalian reproduction is fundamental to higher life on earth. Yet, our understanding of what creates a successful reproduction event is still lacking detail. This in turn hinders optimization of artificial fertilization efforts in both humans and endangered species. The development of sexes, anisogamy and inner fertilization culminated in the appearance of viviparous birth as a trait of nearly all mammalian species in addition to lactation(1). In mammals up to billions of ejaculated sperm are deposited in the female vagina, cervix or uterus(2, 3) and have to travel from the site of semen deposition through the entire female genital tract - which is orders of magnitude larger than the sperm itself - to the site of fertilization. Consequently, the genital tract faces the requirement to assist sperm on its way to the female oocyte and to assure that only the fittest succeed in fertilization. It is a place of sperm selection, conditioning, and storage. Until ultimately one sperm fuses with the female oocyte, a dramatic reduction of sperm number occurs based on stochastic or selective processes. Deep understanding of these processes and the resulting selection gains importance since reproductive success is at risk in humans and in endangered animal species. Between 1973 and 2011 sperm concentration in human semen from unselected men has decreased by 52% (from 99.0 to 47.1 million per ml)(4), approaching the critical sperm concentration considered minimal for native fertilization (20 million sperm per ml)(5). Therefore, assisted reproduction techniques will support human reproduction under recent social conditions(6) and contribute to animal species conservation. However, bypassing natural selection and conditioning of gametes, as required for *in vitro* fertilization (IVF) techniques, can provoke detrimental consequences for the resulting progeny and the species. Many hypotheses on how spermatozoa are guided and selected on their way to the oocyte have been established(7–12), but most details of the anticipated processes remain undescribed. Analyzing these processes helps to assess the significance of the observed decline of human sperm concentration in industrial countries, to optimize sperm preparation for artificial insemination, or to improve sperm selection for IVF. However, experimental options to investigate sperm transit *in vivo* are limited for ethical and practical reasons, and most information stems from *in vivo* studies in farm animals(13), or from sperm behavior *in vitro*. Primary cell culture models and recent microfluidic 3D cell culture models(14) were also applied to study gamete interactions with different parts of the female genital tract recovered from female animals after slaughter or natural death. To understand complex biological systems, predictive mathematical models are required. Describing the process across multiple levels of biological organization (multi-scale) is vital for comprehensive and predictive modelling of a complex process such as the movement and selection of sperm. Here, we analyze sperm transition in the female reproductive tract integrating hypotheses and experimental knowledge with a spatio-temporal multi-scale biophysical model. We established a three-dimensional reconstruction of the female reproductive tract based on implicit functions. This reconstruction serves as environment for an agent-based model (ABM) for sperm migration from vagina to oviduct. In ABM, an agent is a freely moving, decision making entity, which can interact with its environment and other agents. ABM allows to accommodate the sperm properties important for sperm migration. Furthermore, potentially relevant factors as sperm mortality, active motion characteristics, and guidance by rheological and geometrical conditions can be assessed. Experiments on sperm transport in domestic animals revealed that out of the vast number of deposited sperm only some hundreds pass the utero-tubal junction (UTJ)(15, 16), the small connection between uterus horn and oviduct, where fertilization occurs. It takes a few hours to accumulate a sufficient number of sperm in the oviduct to ensure fertilization(13, 15). Our dynamical model for the propagation of sperm to the UTJ predicts key processes, which lead to the reduction in sperm numbers observed *in vivo*. Understanding these processes and shaping constraints of natural fertilization has the potential to greatly improve reproductive success whenever supportive reproduction techniques are required. We first present the mathematical reconstruction of the female genital tract, sperm movement and interaction rules, and environmental conditions affecting sperm migration. The terms sperm and agent are used synonymously. Second, the model is simulated for different scenarios and hypotheses. The bovine reproductive system is used as example since advanced knowledge is available.

## Results

### Spatial multi-scale reconstruction of the bovine female genital tract

The genital tract is considered as a system of connected tubes with variously folded surfaces (Fig. 1, supplementary note 1). We divided it into seven distinct compartments, namely: Vagina, cranial vagina, cervix, uterine body, uterine horns, UTJs and oviducts(1). Three-dimensional cylindrical and conical functions were adapted to mimic their shapes. For instance, the cervix is protruded by primary and secondary longitudinal folds(19), which are mathematically described by trigonometric functions, altering the compartment radius. The individual compartments were combined to an enclosed comprehensive entity by joining them in z-direction. At each compartment transition, e.g. from vagina to cranial vagina, the compartment-describing functions are mathematically identical, ensuring that the system is spatially enclosed.

**Fig. 1.**
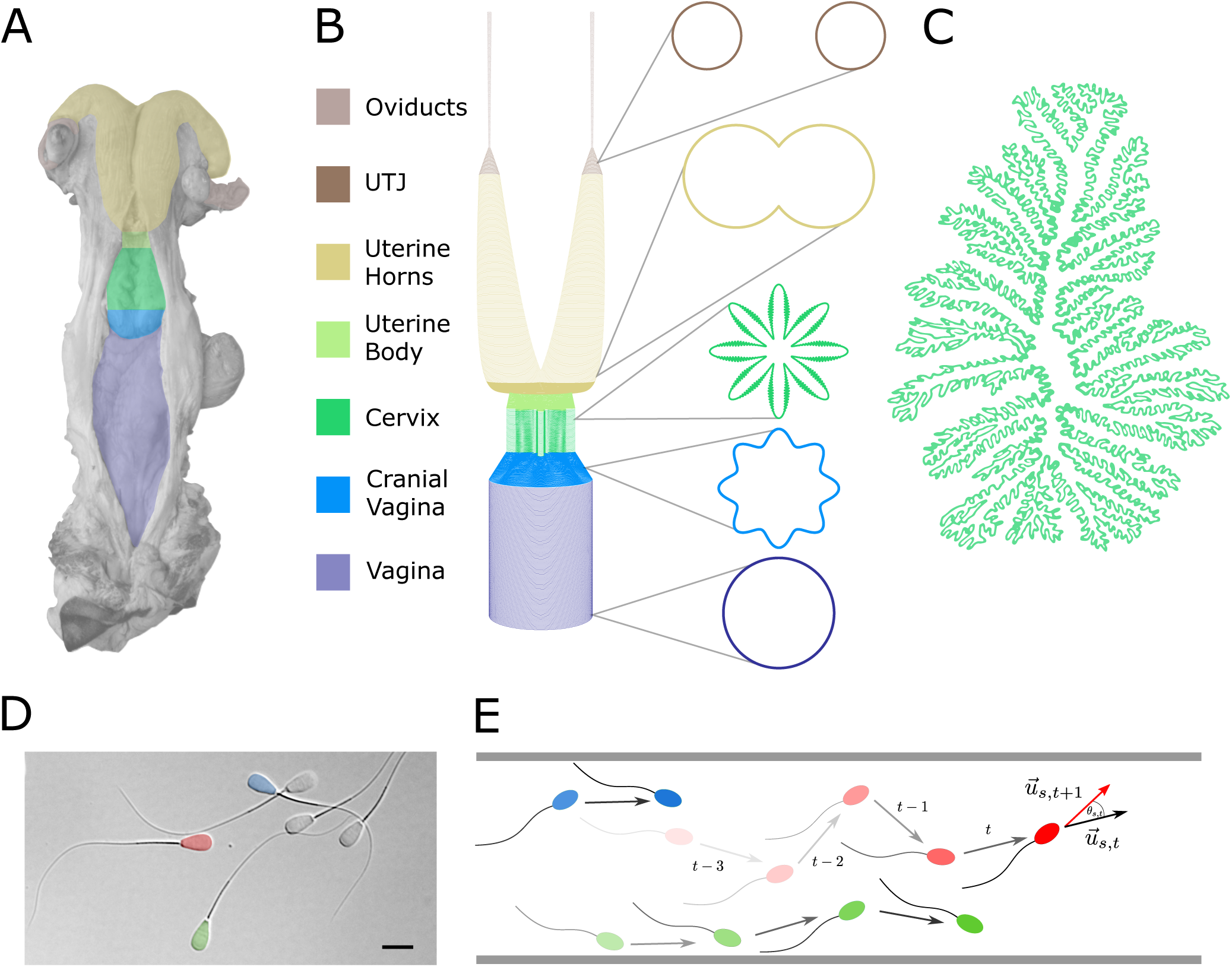
Mathematical representation of the female genital tract and sperm movement. **A** Grey-colored photograph of a bovine female genital tract. Colorcoding indicated separate compartments. The Photograph was kindly provided by Dr. A. Peters IFN Schonöw. **B** The genital tract is mathematically represented by three-dimensional functions sketched here together with cross-sections at different heights. **C** Sketch of a bovine cervical cross section, adapted from (19). **D** Differential interference contrast image of bovine sperm, with colorized heads (to highlight individual differences), bar is 10 μm. **E** Sperms are represented as agents with individual properties in ABM, exploring the reconstructed female tract.

### Temporal model of sperm movement

In the ABM, each sperm cell is represented by an agent *s*. The model is initialized by assigning a random initial position 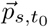 within the cranial vagina to each agent, resembling vaginal deposition(11). The agents have several individual attributes, namely i) average speed 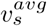, ii) lifetime *τ*_*ls*_, iii) orientation 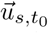, and iv) standard deviation of the deflection angle 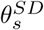 (Tab. 1, Fig. 2). In each time step *δt* a deflection angle *θ*_*s*_ is drawn from a normal distribution around zero with standard deviation 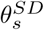 and a current speed *v*_*s,t*_ from a norma ldistribution with mean 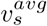 and standard deviation 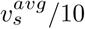. The orientation is updated, by deflecting the orientation vector 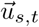 by *θ*_*s*_. To calculate the new agent position, the orientation is scaled by the current speed *v*_*s,t*_:

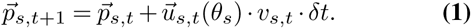

**Fig. 2.**
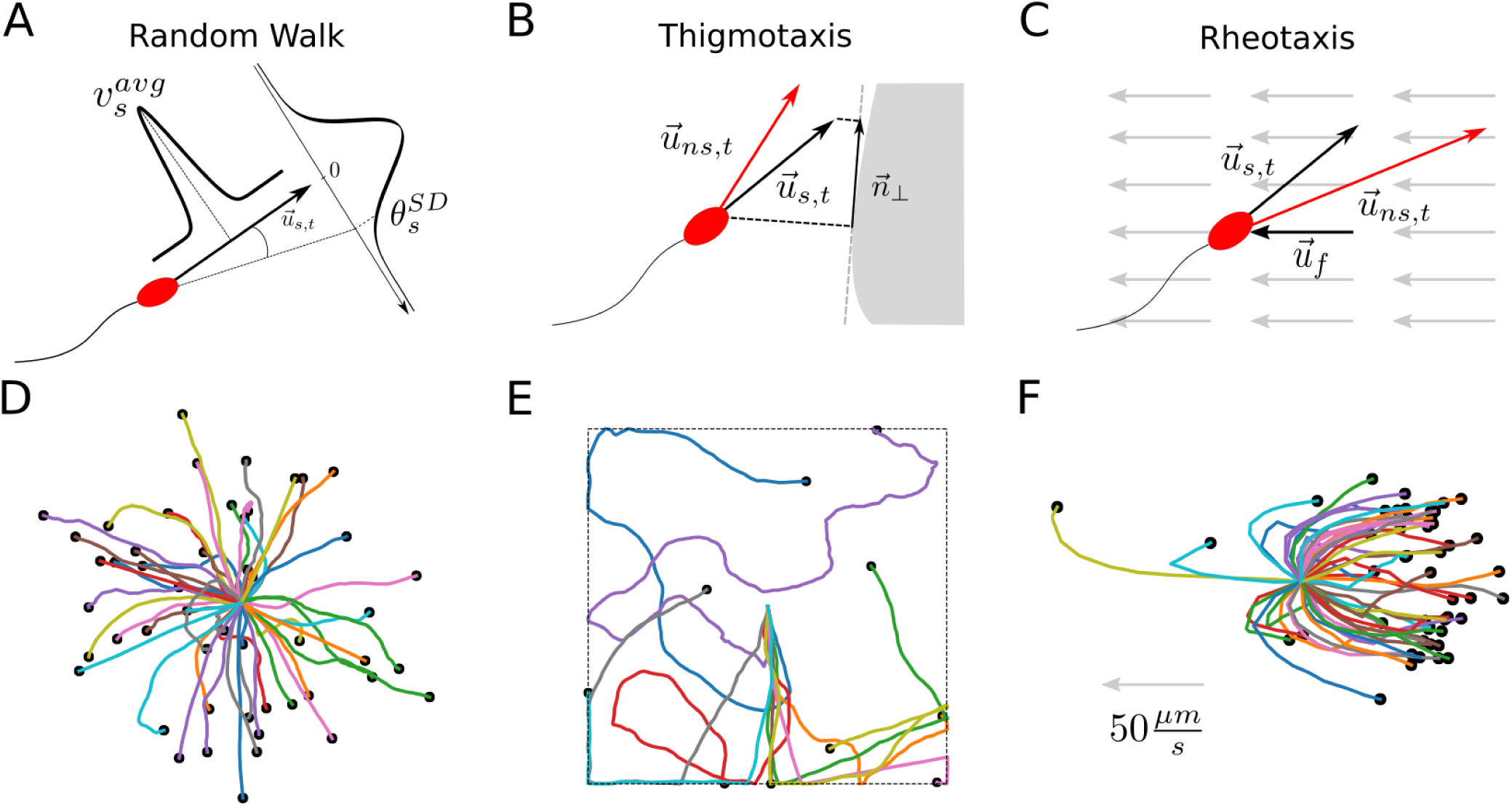
Agent movement, interaction rules, and resulting trajectories in a box. **A** In each time step, agents draw velocities and angles from normal distributions defined by 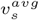 and 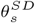. **B** Thigmotaxis is described by averaging the agents orientation vector 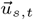 with its projection on the tangent plane of the compartment wall 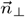, providing the new agent orientation 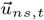. **C** Positive rheotaxis is implemented by subtracting the fluid orientation vector 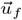 from the sperm orientation 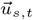 (supplementary note 2). **D-F** Agent movement simulated in a box: agents started in the center, black dots depict final positions. **D** Random walk **E** Thigmotaxis: agent movement restricted by walls (dashed lines). Starting with the same orientation, agents adapt their orientation when hitting walls. **F** Positive rheotaxis: facing a fluid flow of 50 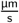 (direction indicated by arrow), agents reoriented into the flow and moved against it.

**Table 1.** Major sperm parameters and their origin. Speed and lifetime were drawn from normal distributions 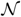. The standard deviation of the deflection angle and the initial orientation were obtained from uniform distributions 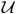.

| Parameter | Distribution | Description | Source |
| --- | --- | --- | --- |
| $v_s^{avg}$ | $\mathcal{N}(60; 10) \frac{\mu\text{m}}{\text{s}}$ | Average velocity | Hyakutake et al.(17) |
| $\tau_{ls}$ | $\mathcal{N}(24; 6)\text{h}$ | Lifetime | estimated |
| $\theta_s^{SD}$ | $\mathcal{U}(1; 119)$ | Standard deviation of deflection angle | Comparison with Tung et al.(18) |
| $\vec{u}_{s,t}$ | $x, y, z \in \mathcal{U}(0; 1)$ | Orientation | |

Thus, sperm movement without any interaction is a spatially restricted random walk.

### Box model reproduces *in vitro* dynamics of sperm

Using results of elaborated sperm cell tracking techniques(20, 21) and a descriptive set of movement parameters as defined within the Computer Assisted Sperm Analysis (CASA)(22), we simulated agent movement in a box with a height of 20 µm representing a typical specimen chamber (Fig. 2) and tested different phenomena as described below. Basic movement and random walk are shown in Fig. 2A,D.

Sperm align to surfaces and edges(23), a process called **thigmotaxis**(24). This behavior is realized by alignment of agents to the compartment boundary, by averaging the agent orientation vector 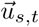 with its projection 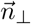 on the approximated tangent plane (Fig. 2B, E, supplementary note 2). The corrected orientation 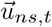 approaches the compartment wall. Hydrodynamically, sperm cells are pushers, pushing fluid onwards and rearwards while replenishing it from the sides(25). As a consequence, sperm aligned to a surface have a lower probability to change direction than free swimming sperm. This is implemented by diminishing the deflection angle with increasing alignment (Supplementary Eq. S30). Mucus is predominantly present in the cervix and potentially guides the sperm towards the uterus and oviducts, as it performs **positive rheotaxis** and orients itself against an oncoming flow(19, 26). Mucus flow was described by a vector field in which its speed increases quadratically with distance to the compartment boundary, resembling a Poiseuille profile. The maximum fluid speed 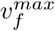 was defined at the lower end of the cervix. Assuming a continuous volume flow through the system, ensuring that no mass vanishes from the system, the mucus velocity *v*_*f*_ at each point in the system was calculated (Fig. S6). The flow direction 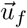 points towards the vagina and is defined compartment-wise (supplementary note 2). Generally, the faster the fluid and the agent, the better the agent reorients against the fluid direction. Thus, sperm orientation 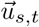 is updated by i) averaging it with the weighted fluid flow direction 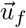 at its position (cross product of 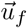 and 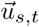) and ii) a logistic term, resulting in faster alignment when sperm swims perpendicular to the flow and when fluid velocity is above a threshold, as sperm only align in sufficient flow rates(18)(Fig. 2C, F, Eq. S46/Eq. S47).

### Cell persistence reproduces experimental data

Knowledge is sparse on the angular deflection of sperm per time. Within CASA, straightness (STR) is calculated by the vector length of displacement divided by the contour length of a sperm trajectory. Tung et al.(18) measured STR values of 0.87 ± 0.02 for bovine sperm tracked for 2.81 s. Using this value we estimated the deflection angles for sperm move>ment, by simulating agent movement in a box and calculating their STR(18, 27). The agents were positioned in the middle of the chamber, restricting movement to z-direction. Calculation of the STR showed that (i) it is timestep independent, which is ensured by the Euler-Maruyama method(28), and (ii) that it perfectly agrees with the measurements from Tung et al(18) (Fig. S9).

### Simulations in the reconstructed female tract

To investigate the success of sperm in the female tract and assess the impact of thigmotaxis, mucus flow, immune system, and STR, we simulated the adjusted ABM within the reconstructed genital tract model. Fig. 3 shows an example of a typical ABM simulation, demonstrating how sperm explore the genital tract over time (see Supplementary Movie 1). For this simulation, a mucus flow (maximal mucus velocity in cervix: 50 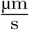) was applied and agents move through the cervix into the uterus cavity. At 0 hpi, sperm-representing agents are confined to the cranial vagina, where they are initialized. Agents either obtaining negative z-position or having successfully reached the oviducts were removed from the simulation, respectively.

**Fig. 3.**
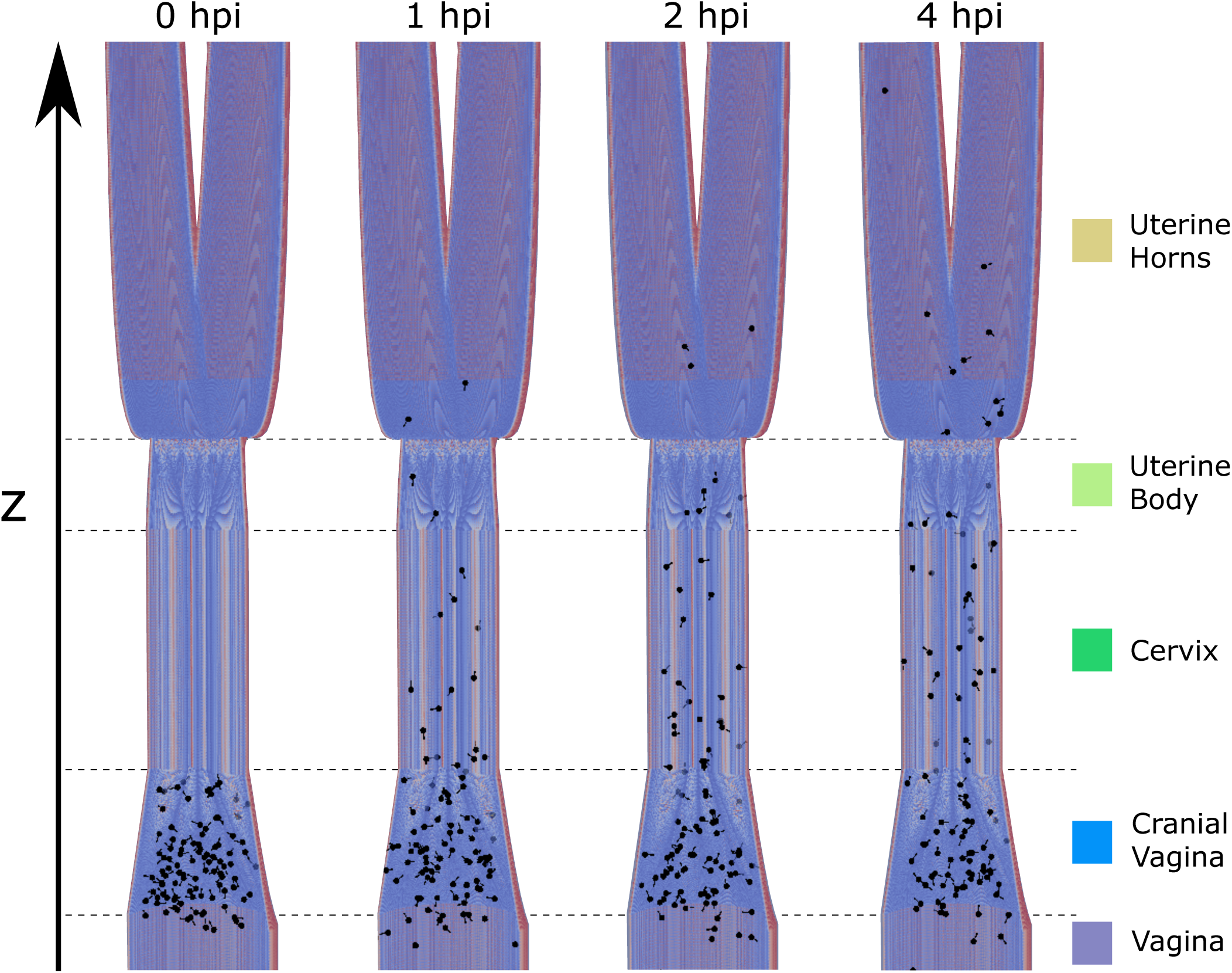
Sperm migration in the female genital tract. Agents are positioned in the cranial vagina and explore the space over time in hpi (hours post insemination). The z-position of the agents is the main observable of the system, defining in which compartment an agent is located. For better visibility, sperm were magnified and the genital tract truncated.

### Survival rate is modulated by immune system

Semen deposition triggers the invasion of neutrophilic granulocytes, which phagocytize sperm in the reproductive tract during the first hours after insemination(7, 29). Agents have a lifetime drawn from a normal distribution (Tab. 1). Immune system activation shortens the agents’ lifetimes. The lifetime decrease is described by a sigmoidal function (Fig. S8). Whenever an agent’s lifetime drops below zero, indicating the sperm’s death, it is removed from the simulation. Mullins and Saacke(19) proposed that sperm is protected from immune cells within the secondary folds (microgrooves). Therefore, being in a microgroove protects agents from lifetime reduction. Thus, the effect of the immune system depends on the agent’s position and the time after insemination and, hence, affects agents differently.

### Wall alignment facilitates directed motion

To test the effect of thigmotaxis alone on the agents’ performance, we simulated scenarios with and without thigmotaxis (Fig. 4). When alignment was omitted, i.e. agents randomly moved through the reproductive tract, none out of 8 million agents reached the oviducts. With thigmotaxis, 336 of 4 million (0.0084%) agents reached the oviducts. Some agents quickly bypass the cervix and move through the uterine horns, while the majority remains within the cervix. After 2 hpi, the first agents reached the oviducts. The agent number drastically decreases between 2.5 and 4 hpi, mostly caused by removal of agents due to the modeled immune system. Around 0.5% of the agent number reduction is caused by agents leaving the system through the vagina.

**Fig. 4.**
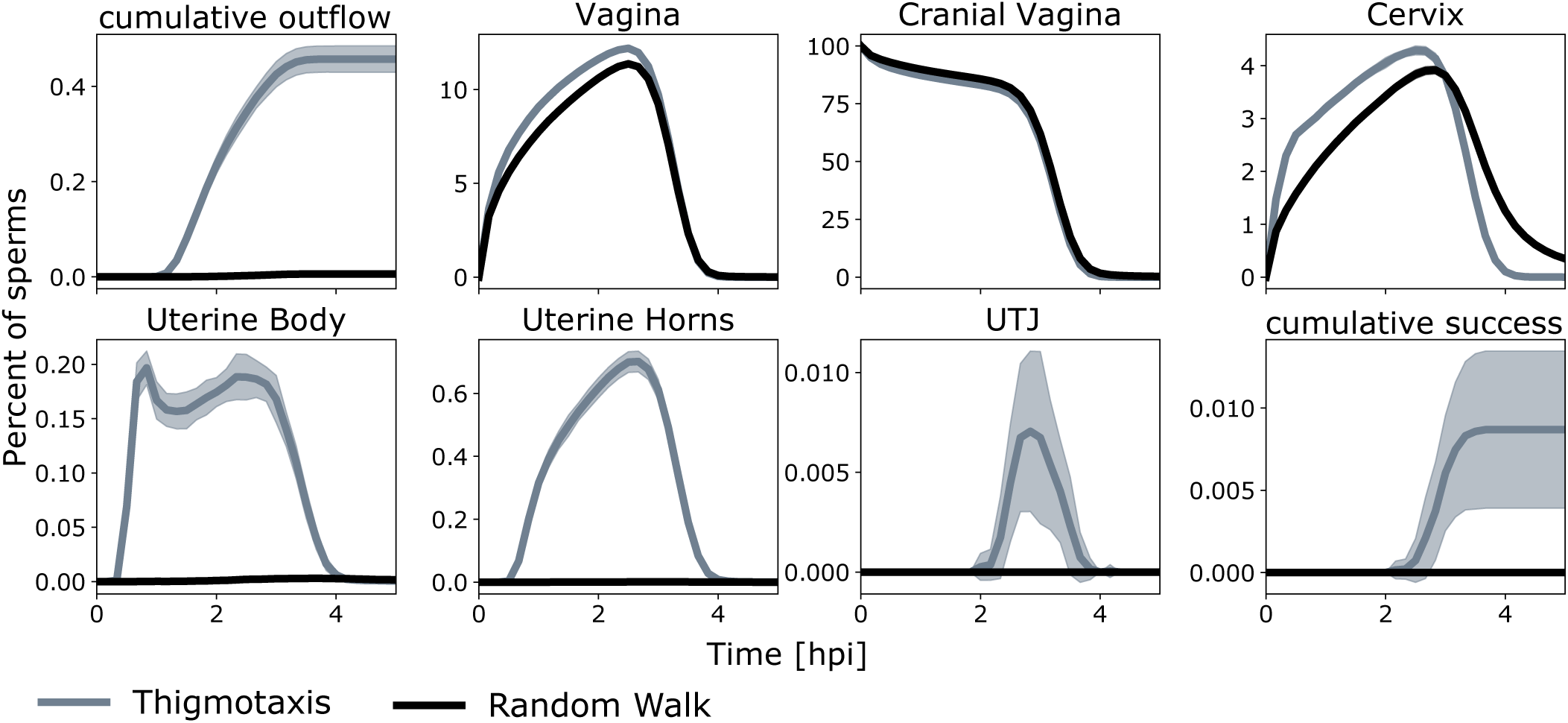
Agent dynamics in different compartments (in %). Upper left and lower right: cumulative time courses of agents being removed from the simulation due to negative z-values (outflow) or reaching the oviducts, respectively. Shaded areas indicate standard deviation. Initially, all agents are located in the cranial vagina. Ultimately, almost all agents were removed by the immune system.

### Fluid flow aligns sperm motion and boosts their success

To evaluate the impact of positive rheotaxis, we simulated different maximal fluid (mucus) velocities. Agents efficiently align into the fluid flow (Fig. 2F). Fig. 5A shows the propagation of the agent population through the compartments for different fluid velocities. With positive rheotaxis, in addition to thigmotaxis, sperm are more likely to leave the cranial vagina towards the cervix. Especially for fluid velocities above 20 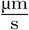, sperm quickly bypass the cervix and begin their transit towards the oviducts. Fig. 5B shows the percentage of successful agents as a function of the maximal fluid velocity. The success rate is maximal for medium fluid flows around 20 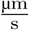.

**Fig. 5.**
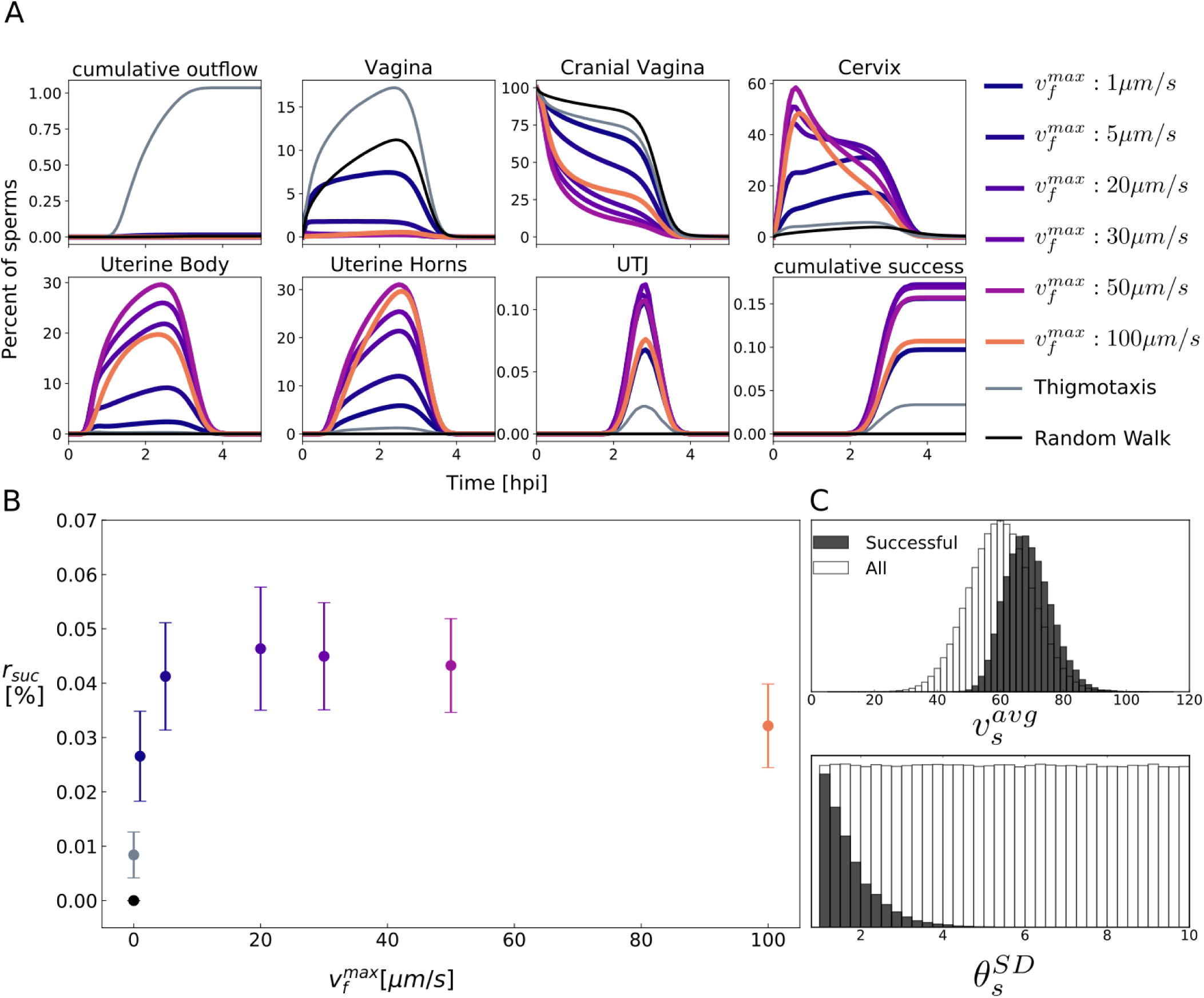
Effect of mucus flow. **A** Percentage of agents per compartment over time. Positive rheotaxis aids agents on their way through the cervix into the uterine body. **B** Success rate of simulated sperm as a function of the maximal fluid velocity. Dots indicate mean percentage of successful sperm, error bars represent the standard deviation. The highest success rate is observed around a maximal fluid speed of 20 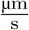. **C** Average speed 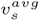 (Successful: 69.9 ± 7.3 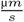, All: 60 ± 10 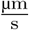) and standard deviation of deflection angle 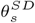 of successful and remaining agents.

### Equiangular swimming increases success probability

Next, we investigated, which sperm properties are important for reaching the oviducts, comparing properties of successful agents with the whole population. It appears that faster as well as equiangular swimming agents have a higher probability to arrive in the oviducts, Fig. 5C. Especially linear movement was a major advantage for the successful agents.

## Discussion

Supporting reproduction is an important aim in human medicine and conservation of endangered species. However, an effective application of assisted reproduction techniques requires detailed knowledge of natural processes ensuring reproductive success. During their transit through the consecutive compartments of the female genital tract, sperm are conditioned for fertilization while their number is dramatically reduced. Conditioning and selection events *in vivo* are far from being understood, but extensive experimental tests are mostly impossible - either due to ethical reasons or due to the lack of animals from endangered species. Therefore, we examined a theoretical approach for its suitability to discover patterns and rules of the journey of sperm to fertilization and developed a spatio-temporal model for the mammalian female reproductive tract. Implementing bovine *in vivo* and *in vitro* data, the model incorporates anatomic reconstruction of and sperm interactions with the female reproductive tract in order to investigate mechanisms of sperm selection and propagation. Despite intensive efforts, it is challenging to make *in situ* observations of propagating sperm within the female genital tract. Spatio-temporal multi-scale modeling of this system enabled us to evaluate the impact of different hypothesized selection processes. For instance, thigmotaxis is well known *in vitro* and it most likely sets a specific environmental condition within the female reproductive tract. Since it remains uncertain whether alignment occurs *in vivo*, we tested the influence of thigmotaxis computationally. In simulations without both thigmotaxis and rheotaxis, no agent reached the oviducts in the given 5-hour-period (Fig. 4), after which all agents were dead. *In vivo*, the first few sperm may reach the UTJ and oviducts even within minutes, and a larger functional sperm population is established until 6 to 8 hours after mating(13). Including thigmotaxis in the simulation leads to a significant amount of successful sperm: By approximating the hydrodynamic properties of sperm, the percentage of agents reaching the oviduct already rose to 0.0084%, which lies in the range of 0.0001% to 0.1% reported by Eisenbach et al.(11) and Reynaud et al.(2), respectively. However, one should be cautious to compare total numbers for mainly two reasons: First, the immune system was modeled solely as a sperm removing process, which is drastically simplified. Second, the deflection angles of sperm were drawn from normal distributions with standard deviations between 1 and 119 degree, which resulted in mean agent straightness values similar to the ones measured by Tung et al.(18). As the real distribution remains elusive and as a small deflection angle is the main characteristic of sperm reaching oviducts (Fig. 5C), a different angle distribution would result in significantly different total numbers of successful sperm. This refers to one of the model’s benefits: to provoke a non-biased re-evaluation of the commonly recorded sperm parameters such as *in vitro* motion characteristics. Further experiments on sperm movement could improve the description of sperm propagation within the model. Considering fluid flow and positive rheotaxis could significantly increase the sperm number reaching oviducts. For a maximal fluid speed of 20 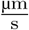, the percentage of agents reaching the oviducts increases to 0.046%. Fluid speeds were measured in mouse oviducts at 18 ± 1.6 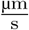 (30). Consequently, the fluid velocity yielding the highest success rate in the model lies in a physiological range. In conclusion, a first model with the potential to describe the entire journey of sperm from insemination to passage of the UTJ was presented and provides estimates on the impact of different selection processes. It revealed that physical properties alone would be sufficient to populate the oviduct with sperm and thigmotaxis turned out to be an indispensable process for successful fertilization. Furthermore, it confirmed fluid flow as a major guidance mechanism, skyrocketing the sperm number reaching the oviduct. In general, it is difficult to distinguish between environmental conditions and non-random processes to assess to what extent sperm reduction on the way to the oviducts is due to stochasticity or selection. Therefore, we performed an analysis of successful sperm compared to the total population. Sperm with the highest velocity and persistence within the implemented distributions had a higher probability to reach the oviducts. Consequently, the model is helpful to guide experimental designs to identify sperm parameters, which had evolved by selection pressure. Sperm successfully undergoing thigmotaxis and rheotaxis should be selected for artificial insemination. In order to answer more detailed questions, more information on genital tract anatomy will be required, e.g. obtaining the exact geometry from medical imaging techniques or pathological sections. This would open exciting possibilities to simulate sperm transport in rare species where the chance to perform *in vivo* investigations is not given at all. The advantage of the presented model is its extensibility. This concerns a more sophisticated simulation of immune responses, inclusion of chemotaxis, thermotaxis, female orgasm as well as sperm interactions with epithelial cells or capacitation-related changes of sperm motion, which primarily occur in the oviduct prior to fertilization. Previous models have already focused either on geometry of the oviducts(31) or on the chemotaxis(32). Therefore, an extended approach modeling the journey of sperm through all compartments will help to discover patterns and rules for a better understanding of selection processes in the context of species-specific reproductive systems and to optimize assisted reproduction techniques such as artificial insemination.

## Methods and Materials

An agent based model for sperm propagation was developed using *Python*. The model initially assigns positions within the cranial vagina compartment to each sperm agent. Furthermore, it sets values for each agent property shown in Tab. 1. Positions and agent properties are represented by arrays, such that the i-th agent was represented by the i-th position of those arrays. Sperm propagation was modelled using update rules for agent position 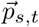, orientation 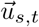 and speed *v*_*s,t*_ in each time step *δt*. Agent movement was restricted to defined shapes, i.e. the reconstructed female genital tract or the box mimicking a specimen chamber. When fluid flow was considered a grid holding fluid velocities was calculated before agent initialization. The spatial resolution of the grid was 20 µm. During simulation agents were exposed to the fluid flow at the nearest point in the grid. Due to the large number of agents, parallelization of the simulation was needed to obtain reasonable simulation times. Parallelization was achieved by usage of a bash script.

### Agent movement

The movement of sperms was described by three rules, (i) by random re-orientation in each timestep, (ii) by alignment along compartment boundaries and (iii) by orientating against the fluid flow. Random re-orientation was implemented as deflection of the agent orientation 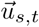 by the deflection angle *θ*_*s*_, followed by a matrix based rotation around the original orientation vector. Alignment along the compartment boundaries was achieved by averaging the agent orientation with its approximated projection on the compartment boundary. Averaging the orientation vector with the negative fluid flow vector described positive rheotaxis. Here, the fluid orientation vector was scaled amongst others by the fluid velocity. More details can be found in the Supplementary Material.

### Data Visualization

Data storage, analysis and visualization was done in *Python*. Export as vtk-format(33) is possible, which can be visualized by *Paraview*(34) (Fig. 3, Supplementary Movie 1).

## ACKNOWLEDGEMENTS

This work was funded by the Deutsche Forschungsgemeinschaft (DFG, German Research Foundation) under Germany’s Excellence Strategy – The Berlin Mathematics Research Center MATH+ (EXC-2046/1, project ID: 390685689 to EK) and IRTG2290 (to EK). Great thanks go to Dr. A. Peters from the IFN Schönow, who provided the Photograph used for Fig. 1.

## Supplementary Note 1: Reconstruction of the bovine female genital tract

All compartments of the female genital tract are considered as connected tubes, each described with a 3D cylindrical or conical function. The different compartments are connected in z direction. All parameters necessary for the description of the female genital tract are given in Table Tab. S1. Fig. S1 shows the result of the described volume and cross-sections at compartment transitions and interesting z positions. How each of the compartments was described mathematically is described below.

### A. Vagina

The vagina is described as a simple cylinder with radius *r*_*v*_, Eq. S1. Length and radius for the bovine vagina were set to *l*_*v*_ = 25 cm and *r*_*v*_ = 2.5 cm, respectively. The cylindrical function *f*_*v*_ for the vagina is given by:

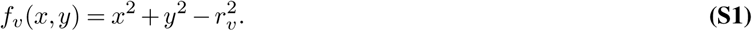

### B. Cranial Vagina

The Cranial vagina compartment mimics the transition from vagina to cervix. Vagina and cervix mainly differ in two ways. First, they have a different radius. Second, the cervix holds primary and secondary folds. Therefore, the cranial vagina compartment must changes in radius and introduces primary and secondary folds dependent on the z-axis. The respective set of equations reads as follows and is explained below:

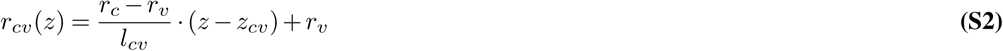

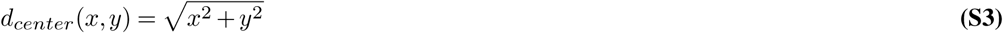

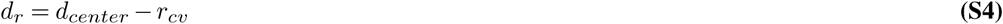

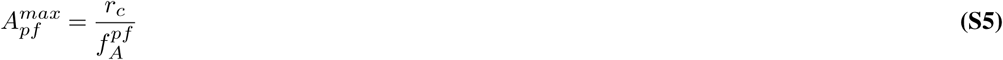

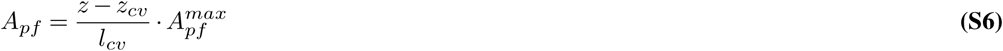

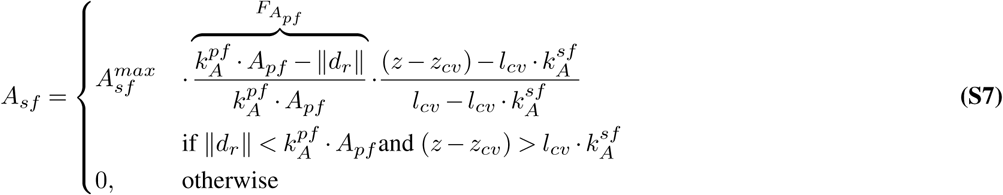

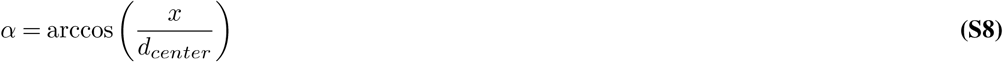

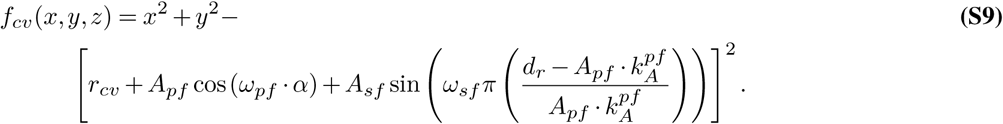

Eq. S2 defines the change of the cranial vagina radius *r*_*cv*_ from the vaginal radius *r*_*v*_ to the cervical radius *r*_*c*_ over the cranial vagina length *l*_*cv*_.

Eq. S3 and Eq. S4 describe the distance to the compartment center *d*_*center*_, that is the z-axis, and the distance to the nearest point on the cranial vagina radius *d*_*r*_, respectively. *A*_*pf*_ (Eq. S6) defines the amplitude (or depth) of the primary folds, which increases linearly with increasing z. Eq. S7 defines the amplitude/depth of the secondary folds *A*_*sf*_. The occurrence of secondary folds is limited in two regards. First, they only occur if primary folds are sufficiently large (which depends on the z-axis), i.e. when the difference between the z-coordinate and *z*_*cv*_ (the compartment baseline) is larger than *l*_*cv*_ times a constant 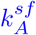, then the secondary fold amplitude *A*_*sf*_ becomes larger than 0. Second, they only occur in the middle of the primary folds, i.e. if the distance *d*_*r*_ is smaller than the depth of the primary fold times a constant 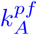 (see first fraction named 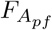 in Eq. S7, depicted in Fig. S2). Here, 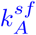 and 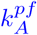 represent the fractions of cranial vagina length and primary folds regions in which secondary folds occur, respectively, while 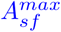 is the maximal depth of secondary folds. Eq. S8 defines the angle towards the x - axis. Finally,Eq. S9 summarizes the contributions for the whole compartment. The squared x and y terms show that the equation still originates from a cylindrical description. The cosine term introduces *ω*_*pf*_ primary folds with amplitude *A*_*pf*_, while the sine term introduces *ω*_*sf*_ secondary folds with amplitude *A*_*sf*_. *ω*_*sf*_ is scaled by *π* and an additional term which ranges between 2 and 0, describing the whole circle, resulting in *ω*_*sf*_ secondary folds per primary fold.

### C. Cervix

The cervix holds primary and secondary folds, but compared to the cranial vagina it has a constant radius. Secondary folds are again limited to regions where primary folds are sufficiently deep, see Eq. S11. The entire cervix is described by Eq. S12. Note that the distance to the radius is defined in the same manner as for the cranial vagina (compare Eq. S4 and Eq. S10), but that the radius used in the compartment equations (Eq. S9 and Eq. S12) is fixed (*r*_*v*_). The equations for the description of the cervix read:

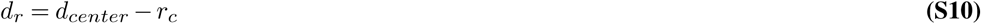

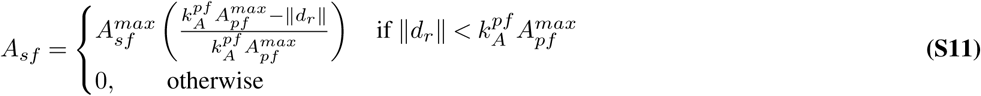

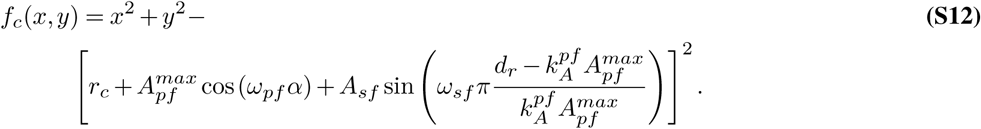

### D. Uterine body

Similar to the cervical vagina compartment, the uterine body compartment has a changing radius and a decreasing depth of primary and secondary folds to zero. The logic behind the equations is the same as for the cranial vagina compartment. Again, the distance to the radius is defined as before, Eq. S14. Here, *r*_*ub*_ is the radius of the uterine body and *r*_*uh*_1_ the radius at the lower end of the uterine horns. *l*_*ub*_ and *z*_*ub*_ are the length and the z baseline of the uterine body, respectively. The respective equations are:

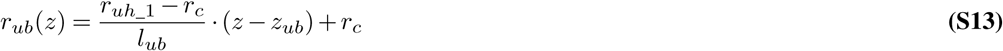

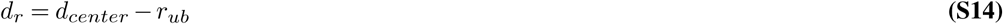

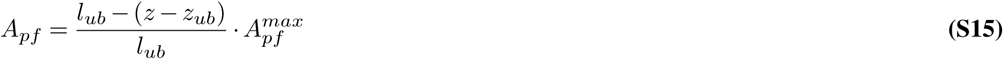

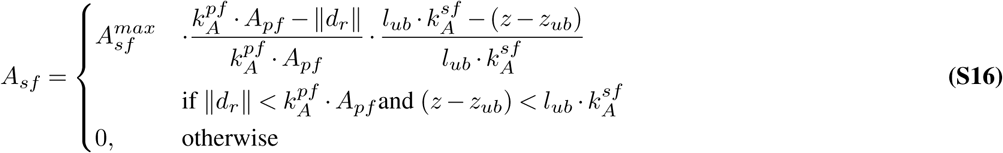

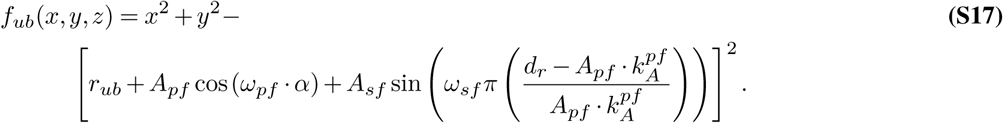

### E. Uterine Horns

The two uterine horns are described by the following equations.

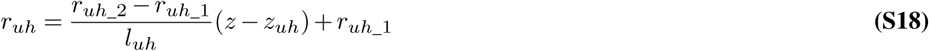

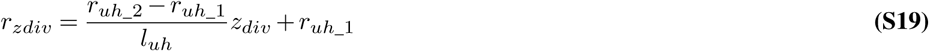

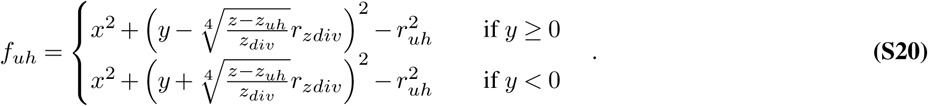

*l*_*uh*_, *r*_*uh*_ and *z*_*uh*_ are length, radius and z offset of the uterine horns, while *r*_*uh*_2_ is the radius at the upper end of the uterine horns. *z*_*div*_ is the z position (with respect to *z*_*uh*_) at which the two horns separate and *r*_*zdiv*_ is the radius of the uterine horns at this height. Eq. S20 defines the shape of the uterine horns, one for *y* ≥ 0 and one for *y* < 0. They have a decreasing radius (Eq. S18) and two centers, which drift apart in the y direction.

### F. Uterotubal junction

In the uterotubal junction (UTJ) the radius decreases again, while the centers don’t drift apart anymore, staying at a constant y position. The transition between the two compartments mimics the narrowest part of the whole tract, with radius *r*_*o*_1_. The equations for the UTJ read as follows:

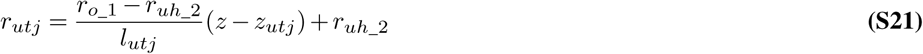

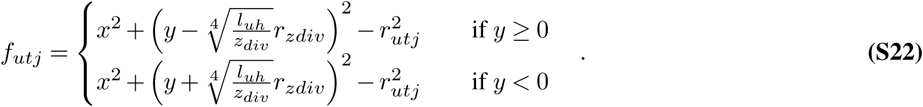

### G. Oviducts

Similar to the uterotubal junction the oviduct compartment only changes the radius of the two parallel tubes (Eq. S23). The end of the oviduct has the radius *r*_*o*_2_. In our simulations this height is never reached, as the question under investigation was when the sperms reach the oviductal compartment. The equations for the oviducts read:

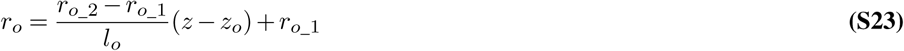

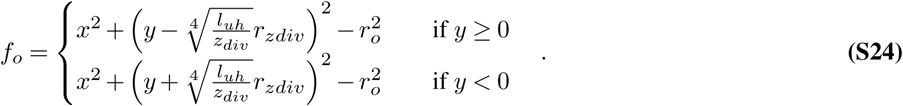

### H. Concatenation

Dependent on the z-position of an agent (described below in supplementary note 2) one of the compartment equations (Eq. S1, Eq. S9, Eq. S12, Eq. S17, Eq. S20, Eq. S22 and Eq. S24) is evaluated. Consequently, the compartments are connected in z-direction creating an enclosed volume. Equation Eq. S25 defines in which case which equation is evaluated.

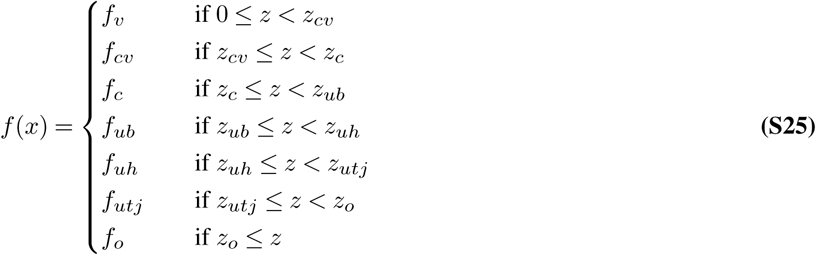

## Supplementary Note 2: Sperm movement

Basic sperm movement is described by a constraint random walk. In each time-step *δt* a new deflection angle *θ*_*s,t*_ and a new speed *v*_*s,t*_ are drawn for each individual agent from normal distributions (Tab. S4). The deflection angle is drawn from a normal distribution with mean 0° and standard deviation 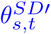. Thus, 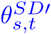 depicts how linear an agent is moving. In the absence of thigmotaxis (supplementary note A) 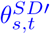 equals 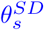.

Each agent gets its individual 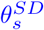 from a uniform distribution, within the limits 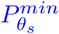 and 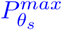, Tab. S2 and Tab. S3. The individual average speed 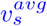 of each agent is drawn from a normal distribution with mean 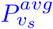 and standard deviation 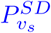, Tab. S2. In each time-step an agent’s orientation 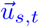 is changes by *θ*_*s,t*_, resulting in a new orientation. Afterwards, the position is updated by equation Eq. S26.

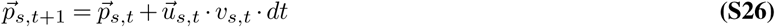

where, 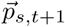 is the updated sperm position. If the new position lies within the volume described in supplementary note 1 the new position is accepted, otherwise the former position is kept. A position lies within the described volume if the function value of *f* ≤ 0 (Eq. S25).

### A. Thigmotaxis

As argued in the main text, sperms can be classified as hydrodynamic pushers, i.e. pushing fluid to the front and to the back, while it is replenished from the sides. Coming close to a wall, the replenishment only takes place from one side and the sperm is pushed and aligned to the wall.

In order to align a sperm’s orientation to the closest compartment wall, the closest point on the wall as well as the wall’s orientation at this point have to be known. This is realized by defining 14 uniformly distributed points around the sperm position. For each of these points it is investigated if it lies inside or outside of the reproductive tract. If a point lies outside of the reproductive tract, the compartment wall is between that point and the sperm. From all vectors which point to the outside, the average vector is calculated and used as the normal vector of a plane. This plane should be approximately parallel to the compartment wall. Afterwards the projection of the sperm orientation on the plane is determined. This projection is used to update the orientation by calculating the average of the projection and the orientation.

The same vectors 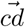, covering all spatial directions, are used for each sperm in order to investigate, whether the sperm is near a compartment wall.

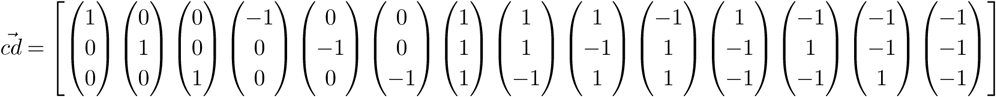

These vectors are normalized and then scaled by (i) their scalar product with the sperm orientation vector and (ii) with half of the length of the sperm. By scaling with the scalar product, vectors which point in a similar direction as the sperm, become longer than those being nearly perpendicular to the sperm orientation. This gives the sperms an ellipsoidal “sensing” zone (Fig. S4), which depends on the sperm’s length *l*_*s*_. Additionally, the length of vectors pointing backwards is reduced to a tenth of their original length, preventing that agents get stuck in corners. Consequently sperms sense walls in front of them earlier than walls on their sides/rear. Also, the length of the sperm plays a role, since longer sperms begin to align earlier. Consequently, each sperm has a set of 14 individual scaled vectors for which it should be checked, whether the sum of each vector with the sperm’s position would lie outside the reproductive tract. The vectors pointing outside the reproductive tract are now averaged and normalized. The resulting vector is used as the normal vector of a plane, which is approximately parallel to the compartment wall. Next the projection of the sperm’s orientation on this plane is calculated (Eq. S27).

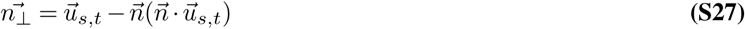

The new sperm orientation is calculated from the former sperm orientation and the projection by Eq. S28.

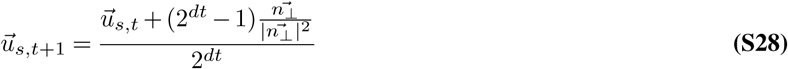

where 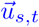 is the sperm orientation and 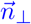 is its projection on the plane. The projection is divided by the square of its length, such that alignment increases the more perpendicular the sperm is to the plane. The term 2^*δt*^ ensures time-step independent alignment, i.e. that the angle by which the orientation is updated is independent of the length of the time-step.

Before the orientation is updated, an alignment score is calculated by the scalar product of the sperm orientation and the projection on the wall, see Eq. S29. This alignment score decreases 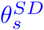 by Eq. S30. As mentioned before without thigmotaxis 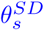 equals 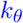, which is the case when *s*_*align*_ = 0.

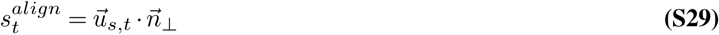

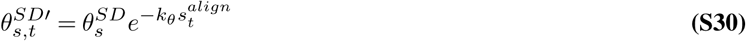

Here, *k*_*θ*_ is a constant defining how strong 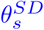 is reduced due to alignment to the surface.

### B. Fluid Field and positive rheotaxis

For the definition of the flow field of fluid within the female genital tract we make four assumptions:

1. No flow through the UTJ and above (oviducts)
2. Flow starting below the UTJ, i.e. in the UTJ-compartment with a fixed flow velocity at the lower end of the cervix
3. A continuous fluid volume profile, with a maximal fluid production in the middle of the cervix
4. A Poiseuille like flow, implying that the velocity increases quadratic with the distance to the compartment boundary

The first point requires that the fluid velocity at the transition between oviducts and UTJ equals 0 (that is a *z*_*o*_). The second point corresponds to the definition of a fluid velocity at the lower end of the cervix (*z*_*c*_). The third point requires the definition of a profile describing the change in volume flowing through the system dependent on the z position in the system, from this the fluid velocity at each point in the system can be calculated. The fourth point requires an expression of the distance to the compartment wall. Here, the distance is the shortest distance to the compartment wall on a certain height.

#### Defining the distance to the compartment boundary

For the circular compartments, i.e. the compartment without folds, the distance is easy to calculate, see Eq. S31.

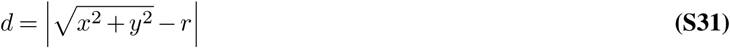

For the other compartments the distance was calculated numerically at pre-defined z-positions. The calculation of the distance was performed in two steps. First, finding equidistant points on the compartment boundary, i.e. on the nullcline of the compartment defining function and second, calculating the minimal distance of every point on the xy-plane to every point on the nullcline. Therefore, we had to define two additional parameters, the resolution in z direction *z*_*sol*_ and the resolution in x and y direction *xy*_*sol*_. In order to find equidistant points on the nullcline, we used the marching squares algorithm implemented in the skimage.measure.find_contours function.

#### Maximal fluid velocity

For two points in the system, we defined the maximal fluid velocity. First, at the transition from the UTJ to the oviduct compartment and second at the transition from the cranial vagina to the cervix compartment. We assume that there is no fluid flux within the transition from oviducts to UTJ. Further, at the transition from cranial vagina to cervix we define a certain velocity, which can be defined for each simulation individually. In order to calculate the fluid velocity at each point in the system we used the recipe below.

1. Calculate the average velocity at the defined z - positions
2. Calculate the volume flow at each of the two z - position (average velocity times area)
3. Use the two volume flow values to define a volume flow profile (here logistic, which maximal change in the middle of the cervix), which defines a continuous volume flow through the entire system
4. Calculate the average fluid velocity in dependence on *z* from the continuous volume profile
5. Calculate the maximal fluid velocity from the average fluid velocity

#### Average velocity at two heights

For the UTJ it is straight forward to calculate the average velocity from the maximal velocity by integrating Eq. S39 over the radius *r*_*o*_1_ and angle *φ* (from 0 to 2*π*), yielding an average velocity defined by Eq. S32:

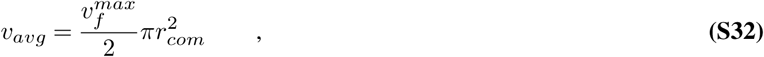

where *v*^*avg*^ is the average fluid velocity at a certain height in the system. For the transition from cranial vagina to cervix the average velocity was calculated numerically, by averaging over the array given by Eq. S40. Next, the average velocities were multiplied by the cross-sectional area at the given heights (*z*_*o*_ and *z*_*c*_), resulting in the volume flow through this cross-section. These two points were afterwards used to define the continuous volume flow profile (see Fig. Fig. S6). Thereafter, the area in dependence of z was calculated. For the vagina, cranial vagina, cervix and uterine body compartment the area was given by 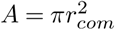. In the case of the uterine horns one has to distinguish between two cases: (1) the area below the height at which two tubes emerge and (2) the area above that point. The area above is simply given by 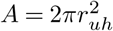, as the uterine horns consist of two circular tubes. For the part of the uterine horns below the division point the cross-section is equal to two overlapping circles and calculated by the following equations:

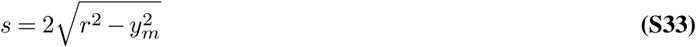

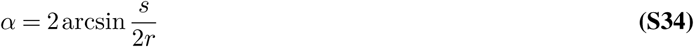

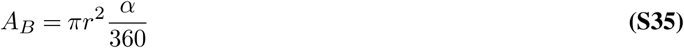

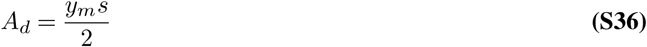

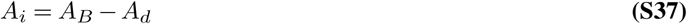

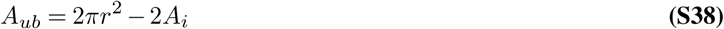

where *s* is the distance between the two contact points of the circle, *α* is the angle spanned by those points and one of the midpoints and *A*_*B*_ is the area of the segment of the circle defined by the midpoint, see Fig. S5. *A*_*i*_ and *A*_*d*_ are the green and orange areas in Fig. S5, respectively.

Dividing the volume flow through the cross-sectional area results in the average fluid velocity in dependence on *z*. The maximal fluid velocity follows from Eq. S32 or by dividing the average velocity by the numerical average of the Poiseuille profile at this *z* position, which corresponds to the average velocity resulting from a maximal velocity of unity. Fig. S6 shows the defined volume flow, the cross-sectional area, the average velocity and finally the maximal velocity in dependence on the z position.

#### Poiseuille Profile

Having the distance to the compartment wall defined we can calculate the fluid velocity at each point by Eq. S39 and Eq. S40, obtaining an analytically or numerically calculated distance respectively.

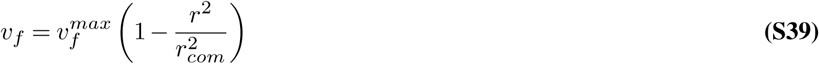

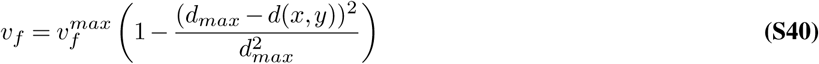

Here, *r*_*com*_ stands for the radius of the system at a certain height, as defined by the different compartment functions. *d*_*max*_ is the maximal distance to the compartment wall and *d*(*x, y*) the distance to the wall at the point (*x, y*). The only unknown entity is the maximal fluid velocity 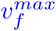. This maximal fluid velocity depends on the z - position and the next paragraph summarizes the underlying assumptions.

Using agent velocities from Hyakutake et al. (17) measured bovine sperm velocities in cervical mucus like medium, it was indirectly assumed that the mucus is a Non-Newtonian fluid. The assumed Poiseuille Profile is valid only for Newtonian fluids. For Non-Newtonian fluids the decrease in fluid velocity towards the boundaries becomes steeper (35), keeping the general appearance of the profile. Therefore the positive effect of positive rheotaxis might even be underestimated.

#### Fluid flow direction

The fluid direction is defined compartment-wise. In the vagina and the cervix, which do not change the radius with height, the fluid is directed down the z-axis. The other compartments are of conical shape and, therefore, the fluid is directed away from or towards the center of the cone, see sketch Fig. S7. For the cranial vagina and the uterine body, the position of the center depends on the primary folds. The center can be defined using the second intercept theorem, Eq. S41 - Eq. S45.

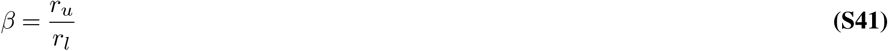

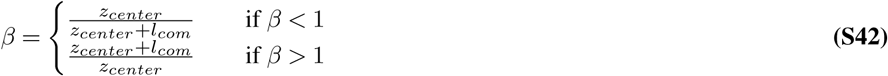

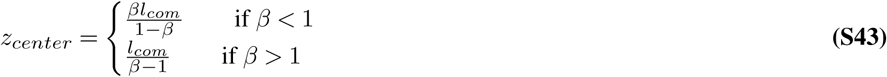

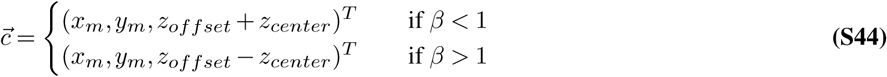

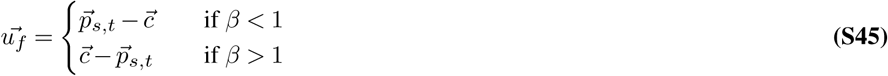

In most cases *x*_*m*_ and *y*_*m*_ are 0. Only in the UTJ compartment *y*_*m*_ is calculated as in Eq. S22. 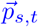 is the position vector of a sperm and 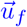 the resulting fluid direction vector, which is normalized.

#### Positive Rheotaxis

As discussed in the main text, sperms align into the fluid flow. Here, we make the following assumptions. First, faster sperms align quicker and second, faster fluid flow leads to faster sperm alignment. The fluid direction vector 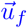 is scaled by Eq. S46 (*sf* for scaling factor). The sperm orientation 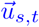 is updated by averaging 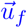 and 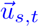 in a time-step independent manner, see Eq. S47.

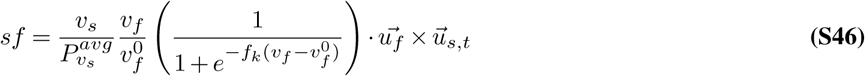

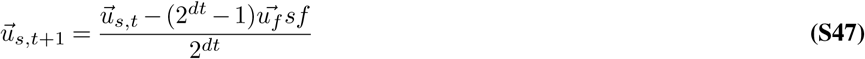

Here, 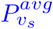 is the average sperm velocity, 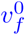 is the fluid velocity at which alignment occurs and *f*_*k*_ is steepness of the alignment response. The logistic term was chosen as it was reported that sperms only align in sufficiently fast fluid flows.

## Supplementary Note 3: Immune system

Each agent possesses an individual life time *τ*_*ls*_, drawn from a normal distribution with mean 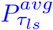 and standard deviation 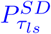 (Tab. S2 and Tab. S3). The immune system is described as a decrease in the life time. The immune system activity is described B Fluid Field and positive rheotaxis by a Hill function with a half maximal activity after 4 hours and a Hill coefficient of 2 (Fig. S8). Given the maximal immune system strength and a default life time of 86400 seconds (1 day) this would be reduced to 30 minutes. Mullins and Saacke proposed that sperms within secondary folds (microgrooves) could be protected from the immune system (19). Therefore agents located in microgrooves do not experience life time reduction by the immune system (Fig. S8).

## Supplementary Note 4: Sperm persistence

Although elaborated sperm tracking techniques exist (20, 21) and a descriptive set of movement parameters is defined within the Computer Assisted Sperm Analysis (CASA) framework (22), knowledge is sparse on the angular deflection of sperm per time. Within the CASA framework, straightness (STR) is calculated by the vector length of displacement divided by the contour length of a sperm trajectory. Tung et al.(18) measured STR values of 0.87 ± 0.02 for bovine sperm tracked for 2.81 s. Using this value we estimated the deflection angles for sperm movement, by simulating agent movement in a box and calculating the persistence (18, 27). With a height of 20 µm, the box represents a typical specimen chamber (22). The agents were positioned in the middle of the chamber, restricting movement to z-direction. Calculation of the straightness showed that (i) it is timestep independent, which is ensured by the Euler-Maruyama method (28), and (ii) that it perfectly agrees with the measurements from Tung et al. (18)(Fig. S9). Sperm agents were simulated in the box, with different time-step lengths *dt* (0.01 s, 0.1 s, 0.5 s and 1 s). We first simulated 50 agents for 2.81 s in order to compare agent persistence with sperm persistence measured by Tung et al.(18)(0.87 ± 0.02). We adjusted the limits of the uniform distribution from which 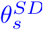 is drawn, such that the persistence of agents were similar to the measured data. Second, we simulated for 3 s with different time steps in order to validate that persistence is time-step independent. Comparison of the simulations with different time-steps showed that the persistence is time-step independent. Fig. S9.

## Supplementary Note 5: Thigmotaxis aids transition through cervix and UTJ

Analysis revealed that wall alignment is especially important while passing the cervix and the UTJ, Fig. S10.

## Supplementary Note 6: Computational Execution

The model was implemented in *Python*. In order to visualize agent movement a *vtk* (Visualization Toolkit (33)) export was implemented, such that the data could be visualized later in Paraview (34). The simulation was parallelized by using a bash script, which started the simulation on different cores.

---

**Table S1.** Parameters used in implicit functions for description of the bovine female genital tract.

| Parameter | Value | Description | Source |
| --- | --- | --- | --- |
| $l_v$ | 25 cm | Length of the vagina | Busch et al.(36) |
| $r_v$ | 2.5 cm | Radius of the vagina | |
| $l_{cv}$ | 5 cm | Length of the cranial vagina | |
| $r_c$ | 1 cm | Radius of the cervix | |
| $l_c$ | 8 cm | Length of the cervix | Busch et al.(36) |
| $l_{ub}$ | 3 cm | Length of the uterine body | Busch et al.(36) |
| $r_{uh\_1}$ | 1.5 cm | Radius at lower end of uterine horns | |
| $r_{uh\_2}$ | 0.5 cm | Radius at the upper end of uterine horns | |
| $l_{uh}$ | 40 cm | Length of the uterine horns | Busch et al.(36) |
| $l_{utj}$ | 5 cm | Length of the UTJ | |
| $l_o$ | 25 cm | Length of oviducts | Busch et al.(36) |
| $r_{o\_1}$ | 0.05 cm | Radius at lower end of oviducts | |
| $r_{o\_2}$ | 0.1 cm | Radius upper end of oviducts | |
| $z_{cv}$ | 25 cm | z baseline of cranial vagina | calculated |
| $z_c$ | 30 cm | z baseline of cervix | calculated |
| $z_{ub}$ | 38 cm | z baseline of uterine body | calculated |
| $z_{uh}$ | 41 cm | z baseline of uterine horns | calculated |
| $z_{utj}$ | 81 cm | z baseline of UTJ | calculated |
| $z_o$ | 86 cm | z baseline of oviducts | calculated |
| $d_{center}$ | - | Distance to z-axis | calculated (Eq. S3) |
| $d_r$ | - | Distance to compartment radius | calculated (Eq. S10) |
| $A_{pf}$ | - | Amplitude of primary folds | calculated (Eq. S6) |
| $A_{pf}^{max}$ | - | Maximal depth of primary folds. | calculated (Eq. S5) |
| $A_{sf}$ | - | Amplitude of secondary folds | calculated (Eq. S7) |
| $A_{sf}^{max}$ | 0.3 | Maximal depth of secondary folds. | educated guess |
| $k_A^{sf}$ | 0.8 | - | educated guess |
| $k_A^{pf}$ | 0.5 | Fraction of primary folds where secondary folds occur | educated guess |
| $\omega_{pf}$ | 8 | Number of primary folds | educated guess |
| $\omega_{sf}$ | 32 | Number of secondary folds | educated guess |
| $z_{div}$ | 4 | Height at which uterine horns divide | educated guess |
| $r_{zdiv}$ | - | Radius at which uterine horns divide | calculated (Eq. S19) |

**Table S2.**
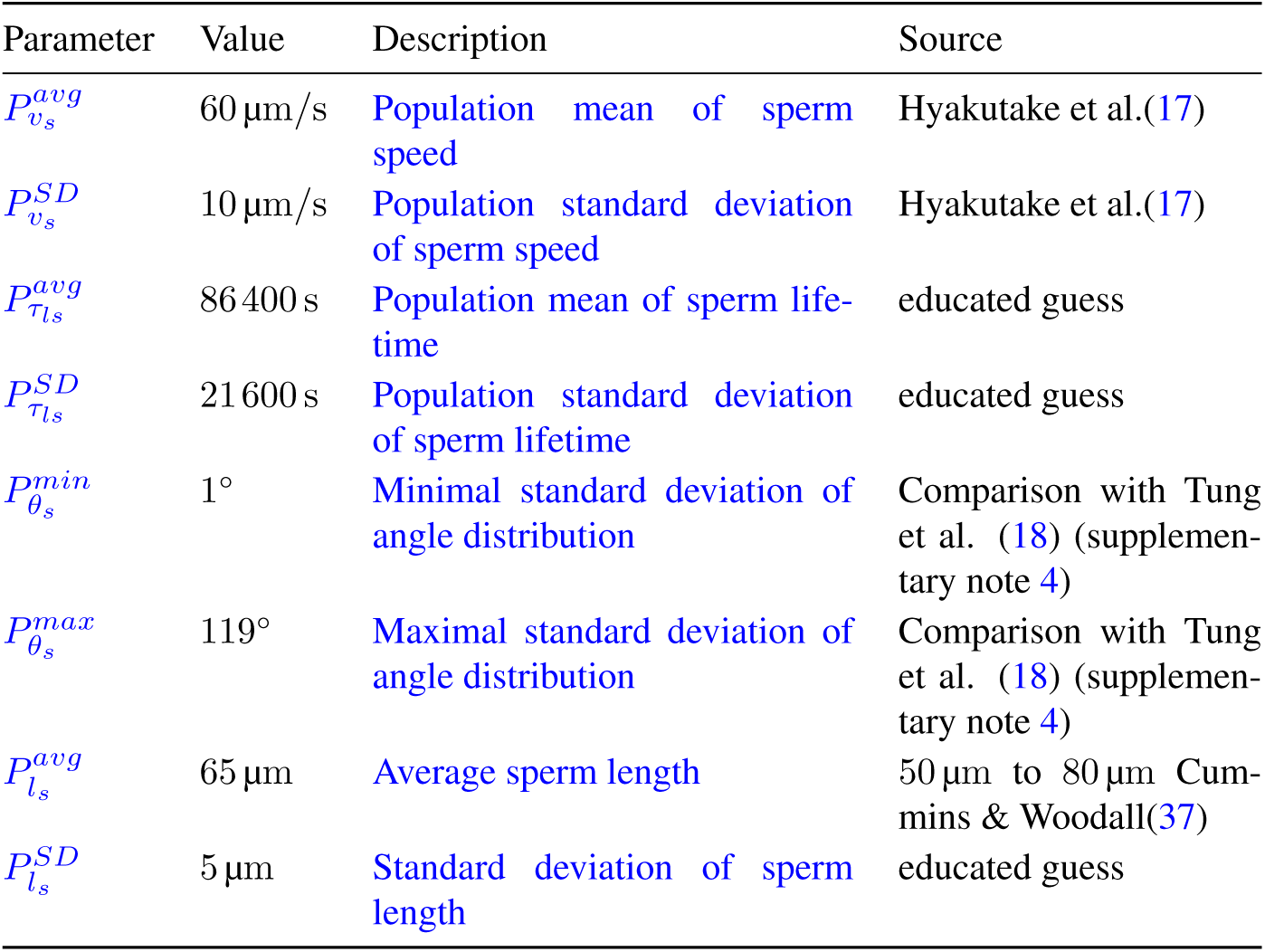
Sperm population parameters.

| Parameter | Value | Description | Source |
| --- | --- | --- | --- |
| $P_{v_s}^{avg}$ | 60 $\mu\text{m/s}$ | Population mean of sperm speed | Hyakutake et al.(17) |
| $P_{v_s}^{SD}$ | 10 $\mu\text{m/s}$ | Population standard deviation of sperm speed | Hyakutake et al.(17) |
| $P_{\tau_{ls}}^{avg}$ | 86 400 s | Population mean of sperm lifetime | educated guess |
| $P_{\tau_{ls}}^{SD}$ | 21 600 s | Population standard deviation of sperm lifetime | educated guess |
| $P_{\theta_s}^{min}$ | 1° | Minimal standard deviation of angle distribution | Comparison with Tung et al. (18) (supplementary note 4) |
| $P_{\theta_s}^{max}$ | 119° | Maximal standard deviation of angle distribution | Comparison with Tung et al. (18) (supplementary note 4) |
| $P_{l_s}^{avg}$ | 65 $\mu\text{m}$ | Average sperm length | 50 $\mu\text{m}$ to 80 $\mu\text{m}$ Cummins & Woodall(37) |
| $P_{l_s}^{SD}$ | 5 $\mu\text{m}$ | Standard deviation of sperm length | educated guess |

**Table S3.** Individual sperm agent parameters.

| Parameter | Distribution |  | Description |
| --- | --- | --- | --- |
| $v_s^{avg}$ | $\mathcal{N}(\mu, \sigma^2)$ | $\mu = P_{v_s}^{avg}, \sigma = P_{v_s}^{avg}$ | Average speed of individual agent |
| $v_s^{SD}$ | - | $v_s^{SD} = v_s^{avg} 10^{-1}$ | Standard deviation of speed of individual agent |
| $\theta_s^{avg}$ | - | 0 | Average angle of individual agent |
| $\theta_s^{SD}$ | $\mathcal{U}(a, b)$ | $a = P_{\theta_s}^{min}, b = P_{\theta_s}^{max}$ | Standard deviation of angle of individual agent |
| $\tau_{ls}$ | $\mathcal{N}(\mu, \sigma^2)$ | $\mu = P_{\tau_{ls}}^{avg}, \sigma = P_{\tau_{ls}}^{SD}$ | Lifetime of individual agent |
| $l_s$ | $\mathcal{N}(\mu, \sigma^2)$ | $\mu = P_{l_s}^{avg}, \sigma = P_{l_s}^{SD}$ | Length of individual agent |

**Table S4.** Temporary sperm agent and other parameters

| Parameter | Origin |  | Description |
| --- | --- | --- | --- |
| $\theta_{s,t}^{SD'}$ | Eq. S30 | - | Diminished standard deviation of individual deflection angle distribution |
| $\theta_{s,t}$ | $\mathcal{N}(\mu, \sigma^2)$ | $\mu = 0, \sigma = \theta_{s,t}^{SD'}$ | Deflection angle |
| $v_{s,t}$ | $\mathcal{N}(\mu, \sigma^2)$ | $\mu = v_s^{avg}, \sigma = v_s^{SD}$ | Temporary sperm agent speed |
| $s_t^{align}$ | Eq. S29 | - | Score for the alignment with a compartment wall |
| $k_\theta$ | 0.5 | - | Strength of thigmotaxis |
| $v_f^0$ | 20 $\mu\text{m/s}$ | - | Sperm alignment velocity |
| $f_k$ | 0.5 | - | Steepness alignment response |

**Fig. S1.**
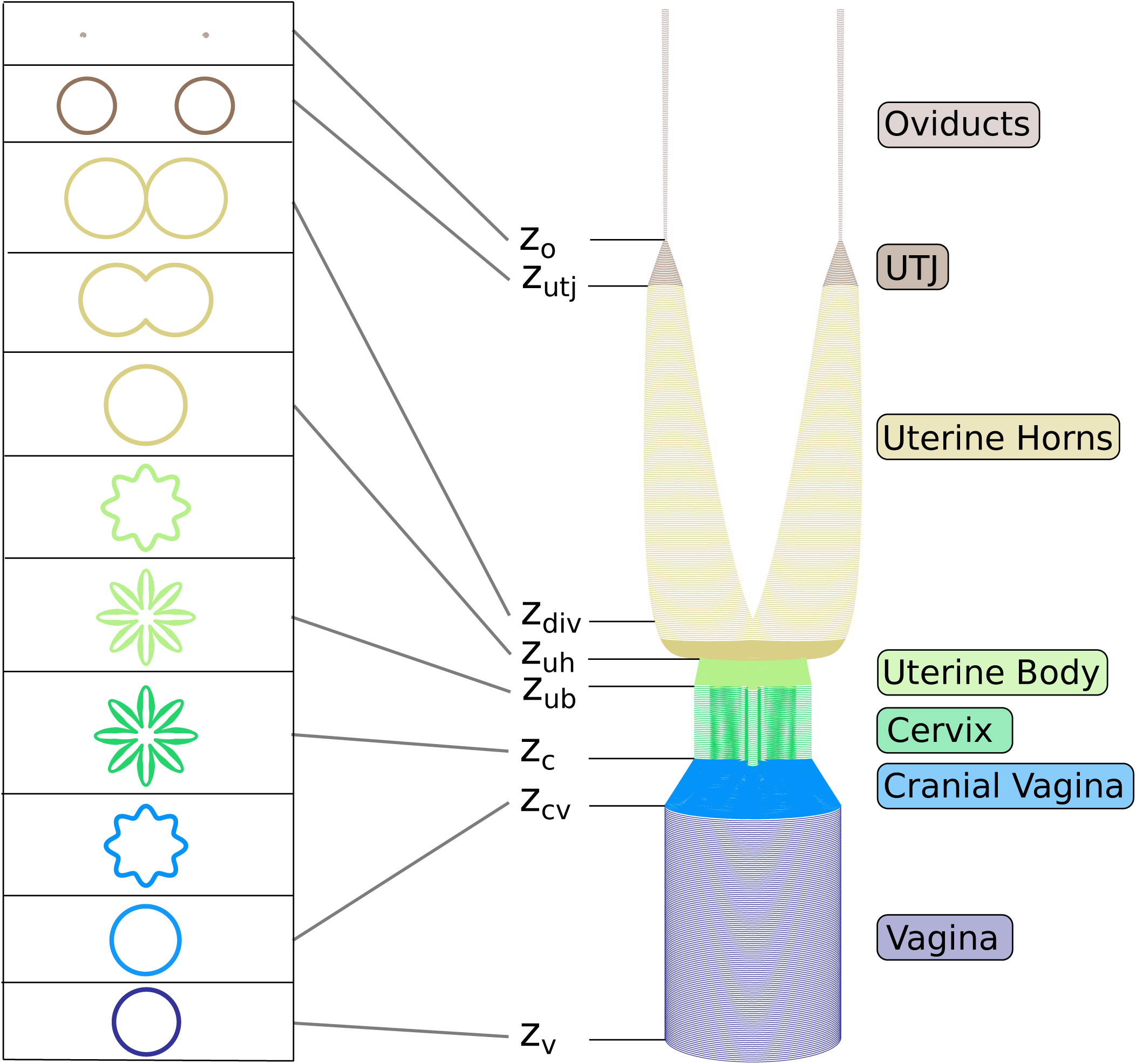
Reconstruction of the female bovine genital tract. Individual compartments (listed on the right) were connected in z-direction. Labelled z-positions indicate compartment transitions and the z-position at which uterine horns divide (see supplementary note E). Cross-sections at these and intermediate positions are shown on the left.

**Fig. S2.**
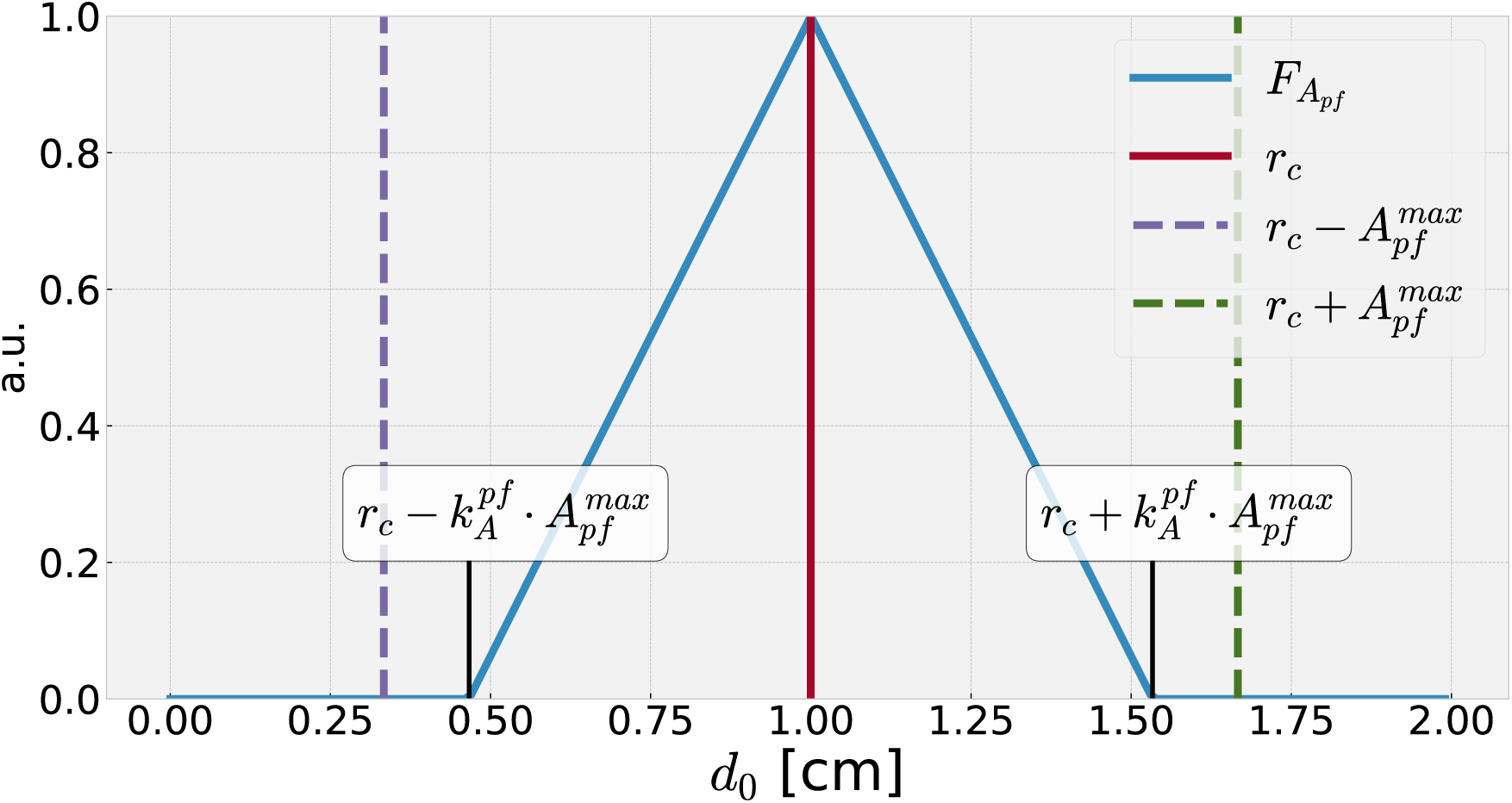
Relative amplitude of secondary fold depth as function of the distance to the primary fold midst. Dashed purple and green line indicate the beginning and end of a primary fold respectively. Red line indicates the center of the primary fold, while the blue line indicates the relative scaling of the secondary fold depth.

**Fig. S3.**
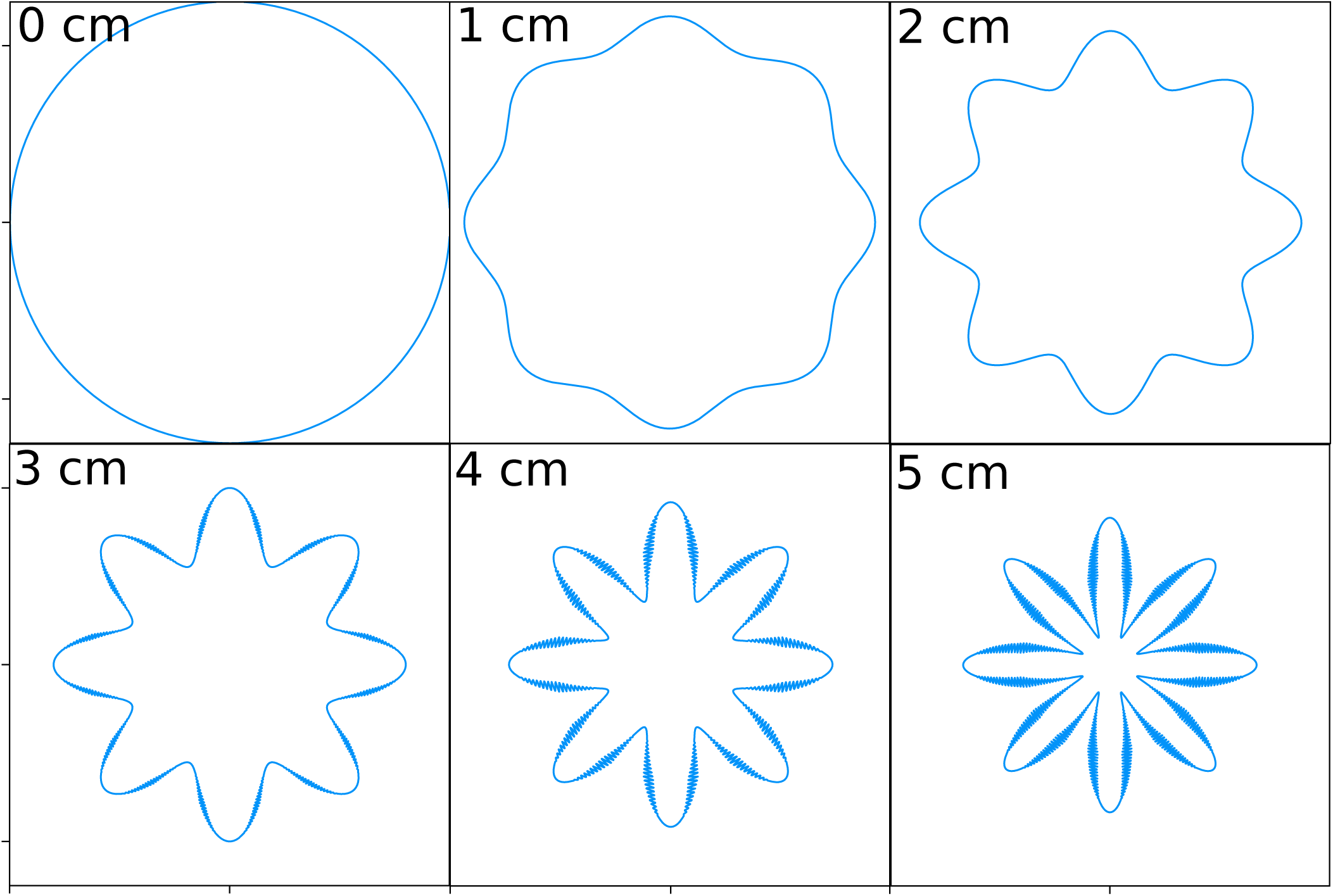
Cross-section of the cranial vagina at different heights measured fromthe vagina. At 0 cm the cranial vagina equals the shape of the vagina, thus the cross-section is a simple circle. With increasing height first the primary (1 cm and 2 cm) and later also the secondary (3 cm, 4 cm and 5 cm) folds develop. Notice that the secondary folds only occur within the upper half of the cranial vagina (restricted in the condition of Eq. S7 by 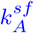) and only in the center of the primary folds (restricted by 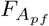). At 5 cm the cross-section equals the cross-section of the cervix.

**Fig. S4.**
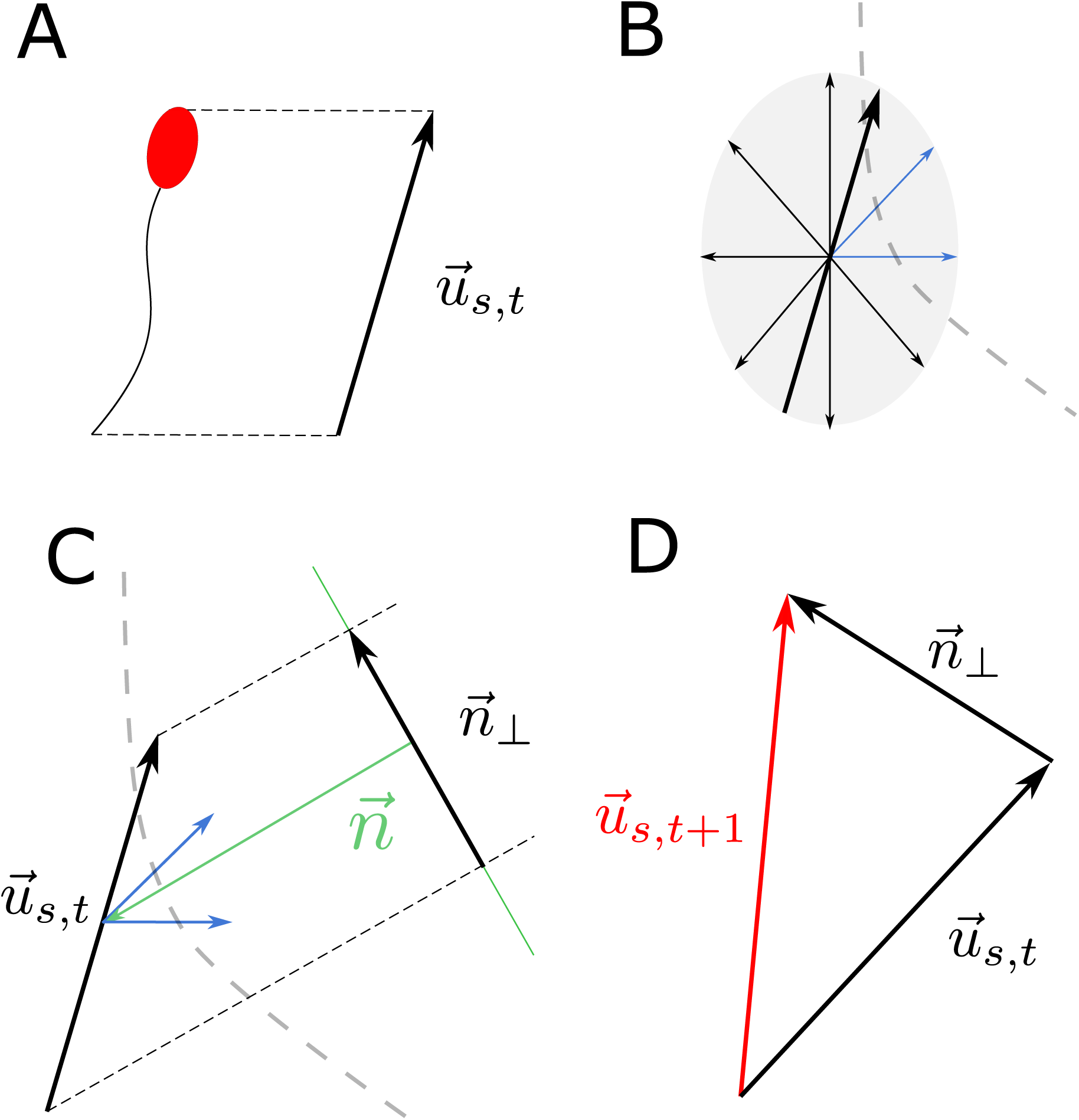
**A** Sperm orientation is defined by its orientation vector. **B** The solid black vector indicates sperm orientation. Smaller black and blue arrows indicate the scaled vectors, for which it is checked if they lie inside or outside the reproductive tract. The scaling provokes an ellipsoidal shape, indicated by the light grey shaded ellipse. The dashed grey line mimics a compartment wall. The two blue colored arrows, point outside the compartment. **C** The weighted average of the vectors pointing outside (shown in blue) of the compartment defines the normal vector of a plane. Sperm orientation as solid black arrow with label 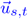. Green arrow indicates resulting normal vector 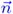. For representational reason it was inverted and enlarged. This normal vector described a plane, shown in dark green. 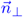 depicts the projection from 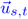 onto the plane defined by 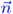. **D** The new sperm orientation 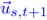 vector is shown in red. It results from the sum of the former direction vector and the projection onto the plane. Subsequent the new orientation vector is normalized.

**Fig. S5.**
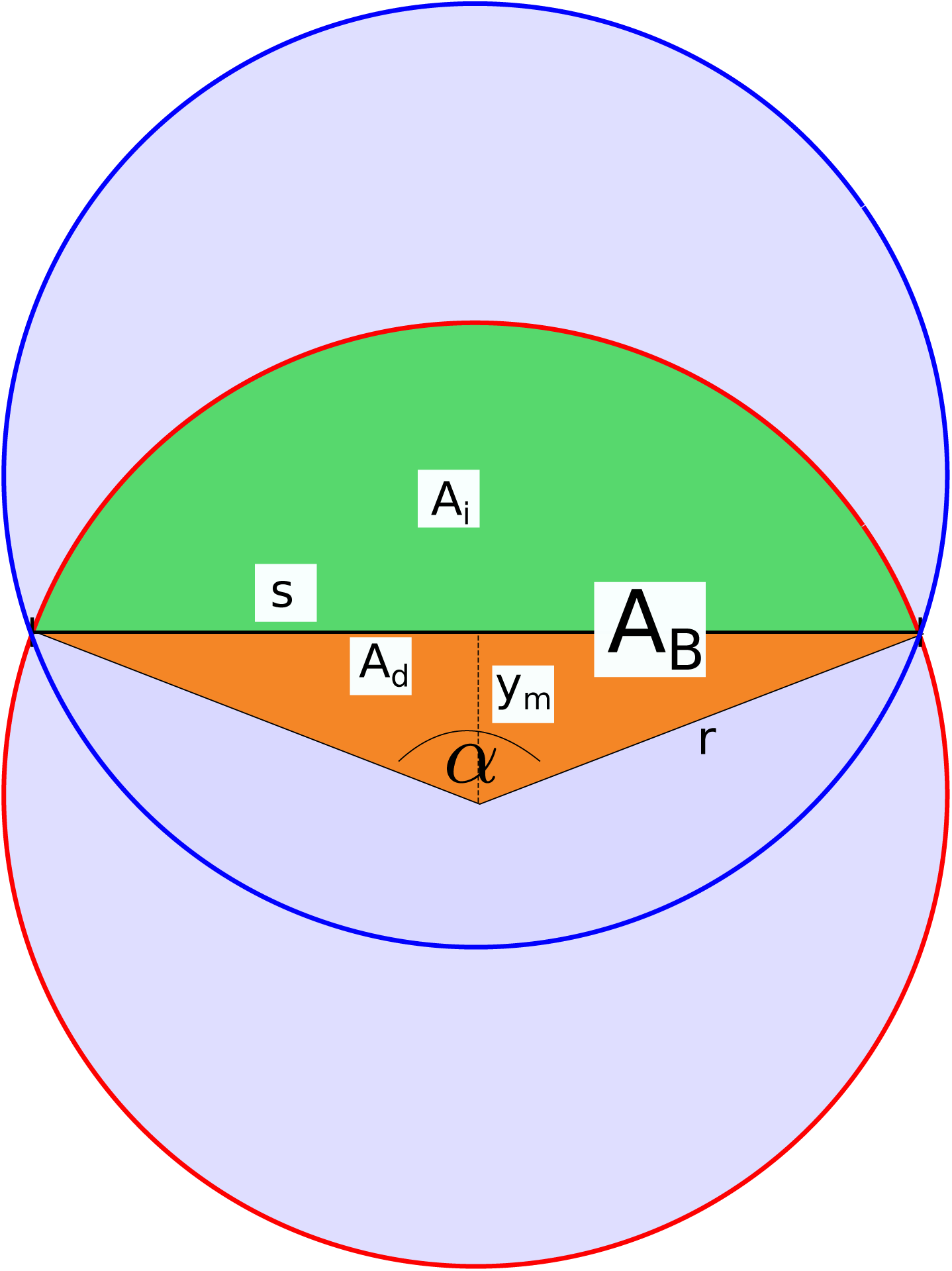
Depicted area calculation for two overlapping circles. *s* is the distance between the overlapping points. *A*_*i*_ is the green and *A*_*d*_ the orange area, while *A*_*B*_ is the area covered by the radiant *α*. *y*_*m*_ is the distance of one midpoint to the line *s*. Values calculated by Eq. S33 - Eq. S38.

**Fig. S6.**
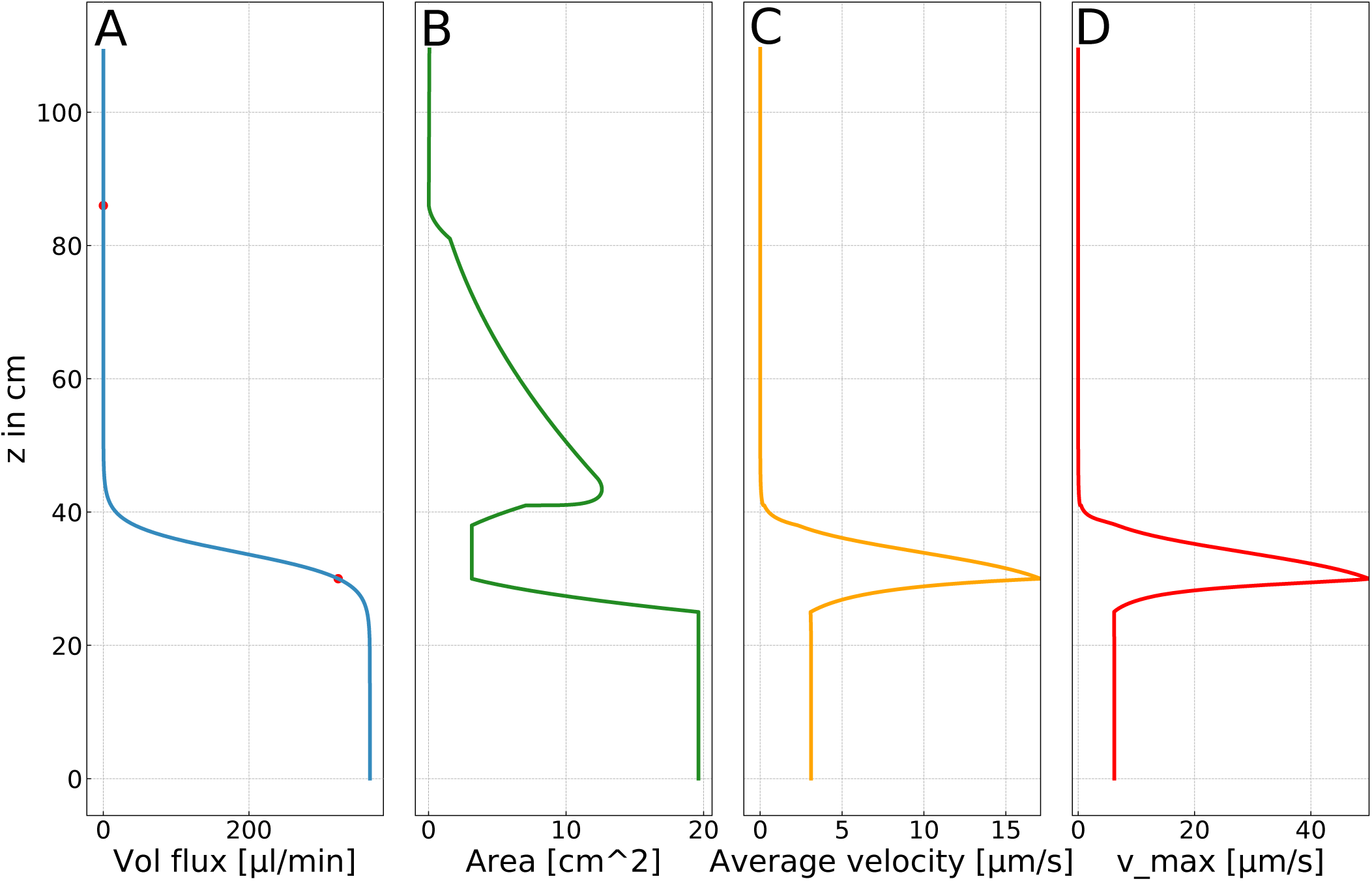
**A** Volume flow. Red dots indicate heights at which the maximal fluid velocity was set. Blue line corresponds to the continuous volume gain throughout the system. The rate of change is maximal in the middle of the cervix compartment (*z* = 34 cm); **B** Crosssectional area as function of *z*. **C** Average fluid velocity calculated from continuous volume flux (A) and cross-sectional are (B). **D** Maximal fluid velocity as function of *z*, calculated from the average fluid velocity (C).

**Fig. S7.**
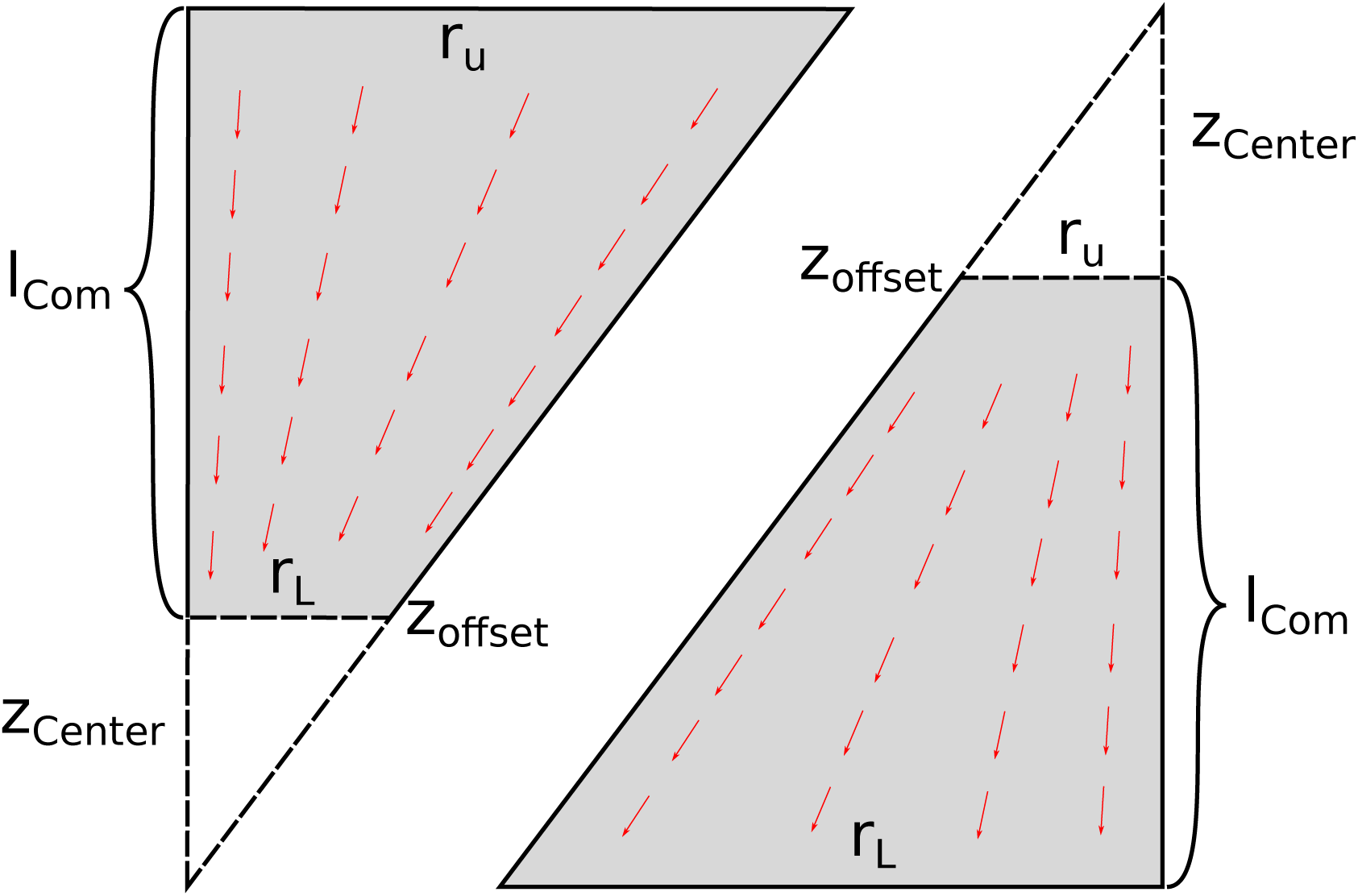
Sketch on how to determine the center of a conical compartment. The compartment is depicted by the grey area. *r*_*l*_ and *r*_*u*_ are the lower and upper compartment radii and *l*_*com*_ the compartment length. *z*_*center*_ is the distance from the cone center to the compartment boundary *z*_*of fset*_. One has to distinguish between the cases that the upper radius is larger than the lower radius (*β* > 1) and vice versa (*β* < 1). In the first case, the fluid flow is directed towards the cone center and in the second case away from the center as depicted by the red arrows.

**Fig. S8.**
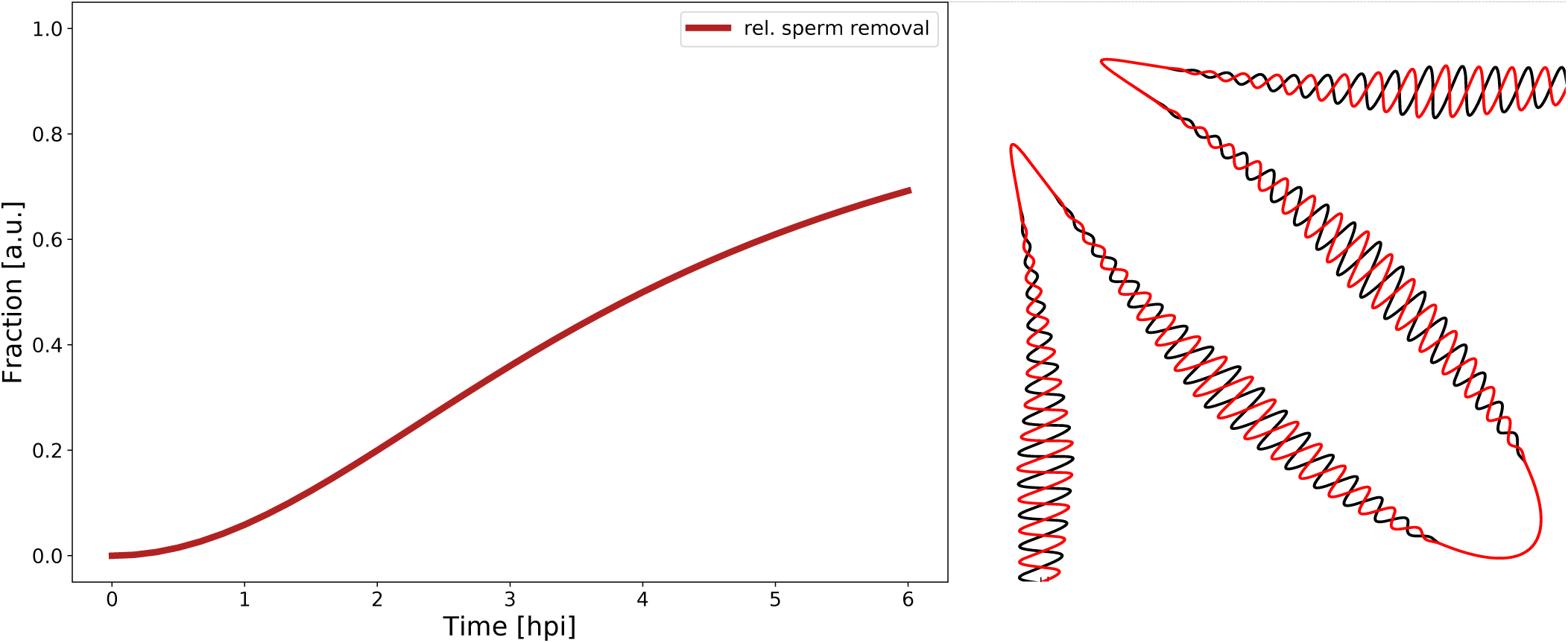
Left: Hill function describing the relative activity of the immune system. Right: Agents within microgrooves are protected from the immune system. An agent is defined to be in a microgroove when it is positioned within the modeled genital tract and outside the female genital tract with inverted secondary folds 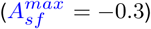. The figure shows an excerpt of the cervical cross-section of the female genital tract with original (black) and inverted (red) secondary folds. Shaded areas depict the cross-section of microgrooves.

**Fig. S9.**
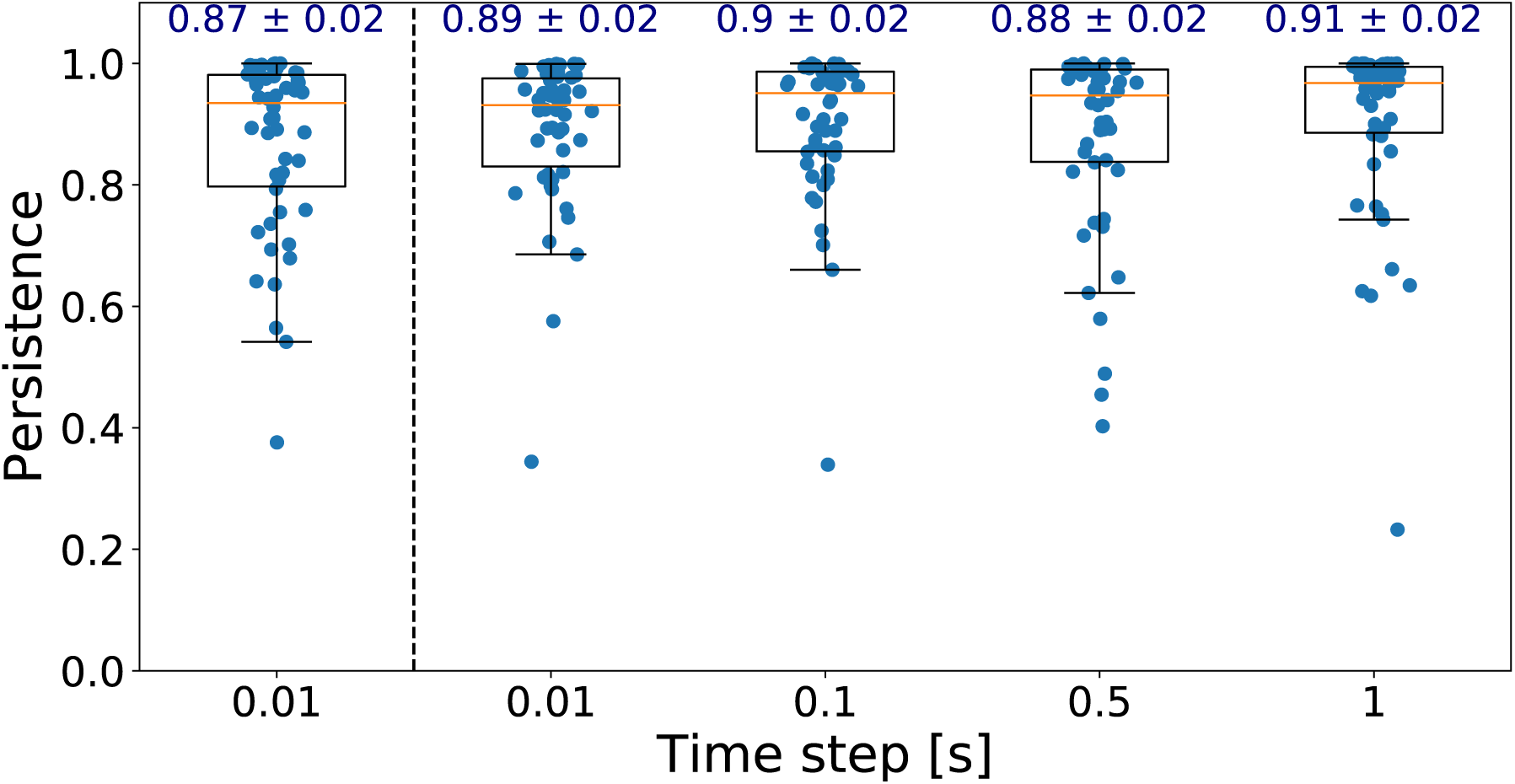
Persistence of simulated agents in a box of 20 µm height. Text above the boxplots gives Mean ± SEM. Orange line indicates median persistence. The persistence shown on the left hand side of the figure originates from a simulation for 2.81 s. For the other persistence the time step was altered and the simulation time was set to 3 s, in order to make persistence comparable between the simulations. Persistence of simulated sperms is in agreement with the persistence reported by Tung et al. (0.87 ± 0.02)

**Fig. S10.**
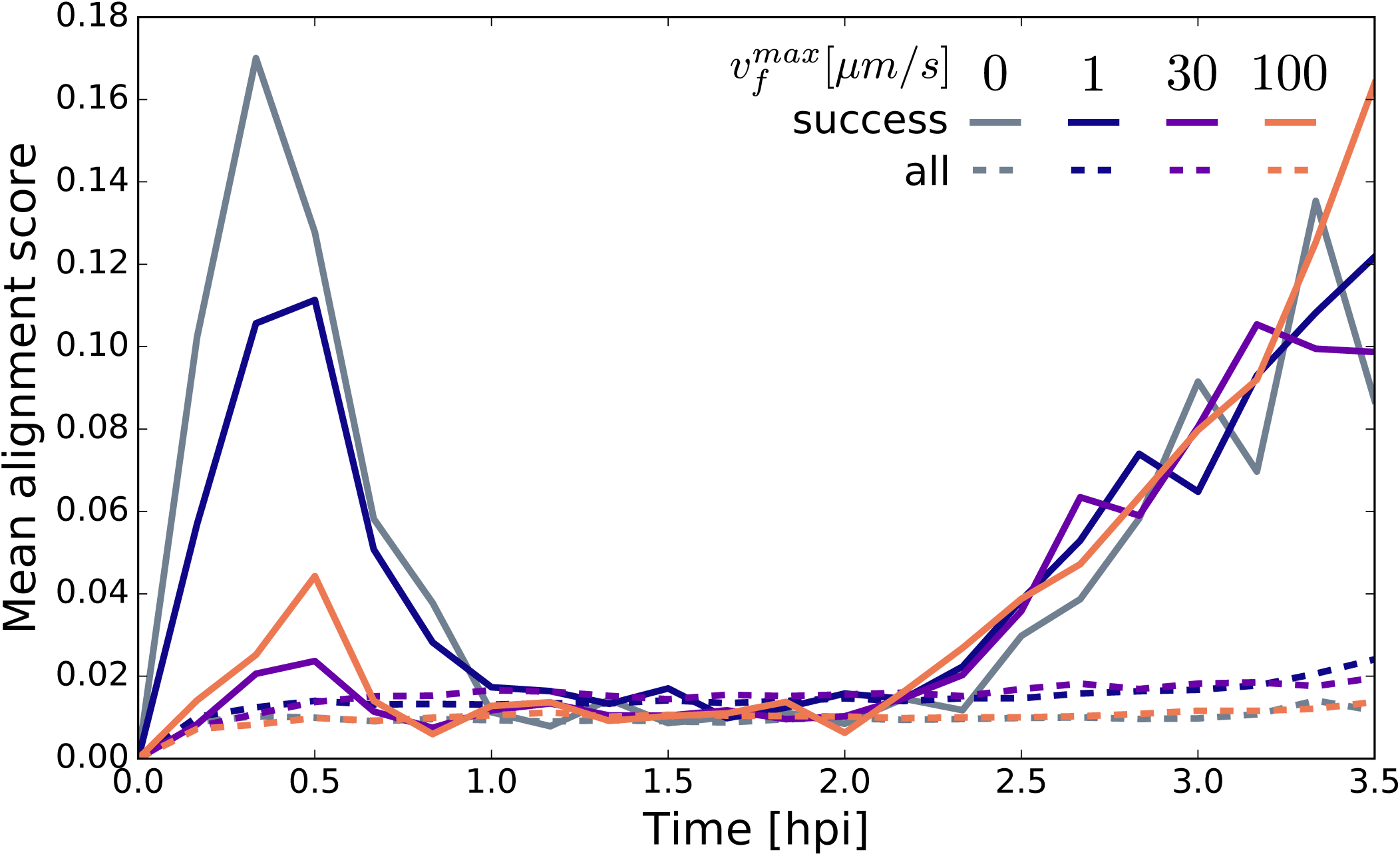
Mean of alignment score 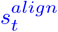 over time. Successful sperms are compared to all sperms. Especially for simulations with only thigmotaxis 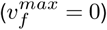 alignment to the wall within the first 30 minutes (while passing the cervix) increases the possibility to be successful. Independent of 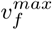 alignment aids the transit through the UTJ (increased mean alignment after 2 hpi.)

## Bibliography

1. Patrick Abbot and Antonis Rokas. Mammalian pregnancy. Current Biology, 27(4):R127– R128, 2017. ISSN 09609822. doi: 10.1016/j.cub.2016.10.046.

2. Karine Reynaud, Zeev Schuss, Nathalie Rouach, and David Holcman. Why so many sperm cells? Communicative & Integrative Biology, 8(3):e1017156, 2015. ISSN 1942-0889. doi: 10.1080/19420889.2015.1017156.

3. Tony M. Plant, Anthony J. Zeleznik, David F. Albertini, Robert L. Goodman, Allan E. Herbi-son, Margaret M. McCarthy, Louis J. Muglia, and Jo Anne S. Richards. Knobil and Neill’s Physiology of Reproduction: Two-Volume Set, volume 1–2. Elsevier Inc., dec 2014. ISBN 9780123977694. doi: 10.1016/C2011-1-07288-0.

4. Hagai Levine, Niels Jørgensen, Anderson Martino-Andrade, Jaime Mendiola, Dan Weksler-Derri, Irina Mindlis, Rachel Pinotti, and Shanna H. Swan. Temporal trends in sperm count: A systematic review and meta-regression analysis. Human Reproduction Update, 23(6): 646–659, nov 2017. ISSN 14602369. doi: 10.1093/humupd/dmx022.

5. Trevor G. Cooper, Elizabeth Noonan, Sigrid von Eckardstein, Jacques Auger, H. W. Gordon Baker, Hermann M. Behre, Trine B. Haugen, Thinus Kruger, Christina Wang, Michael T. Mbizvo, and Kirsten M. Vogelsong. World Health Organization reference values for human semen characteristics. Human Reproduction Update, 16(3):231–245, may 2009. ISSN 13554786. doi: 10.1093/humupd/dmp048.

6. C. Calhaz-Jorge, C. De Geyter, M. S. Kupka, J. De Mouzon, K. Erb, E. Mocanu, T. Motrenko, G. Scaravelli, C. Wyns, and V. Goossens. Assisted reproductive technology in Europe, 2013: Results generated from European registers by ESHRE. Human Reproduction, 32 (10):1957–1973, mar 2017. ISSN 14602350. doi: 10.1093/humrep/dex264.

7. S. S. Suarez and A. A. Pacey. Sperm transport in the female reproductive tract, jan 2006. ISSN 13554786.

8. William V. Holt and Alireza Fazeli. Do sperm possess a molecular passport? Mechanistic insights into sperm selection in the female reproductive tract. Molecular Human Reproduction, 21(6):491–501, jun 2015. ISSN 14602407. doi: 10.1093/molehr/gav012.

9. W. V. Holt and A. Fazeli. Sperm selection in the female mammalian reproductive tract. Focus on the oviduct: Hypotheses, mechanisms, and new opportunities. Theriogenology, 85(1): 105–112, 2016. ISSN 0093691X. doi: 10.1016/j.theriogenology.2015.07.019.

10. John L. Fitzpatrick and Stefan Lüpold. Sexual selection and the evolution of sperm quality. Molecular Human Reproduction, 20(12):1180–1189, dec 2014. ISSN 14602407. doi: 10.1093/molehr/gau067.

11. Michael Eisenbach and Laura C. Giojalas. Sperm guidance in mammals - An unpaved road to the egg. Nature Reviews Molecular Cell Biology, 7(4):276–285, apr 2006. ISSN 14710072. doi: 10.1038/nrm1893.

12. Francisco Alberto García-Vázquez, Iván Hernández-Caravaca, Carmen Matás, Cristina Soriano-Úbeda, Silvia Abril-Sánchez, and María José Izquierdo-Rico. Morphological study of boar sperm during their passage through the female genital tract. The Journal of reproduction and development, 61(5):407–13, 2015. ISSN 1348-4400. doi: 10.1262/jrd.2014-170.

13. R H Hunter and I Wilmut. Sperm transport in the cow: peri-ovulatory redistribution of viable cells within the oviduct. Reproduction, nutrition, development, 24(5A):597–608, 1984. ISSN 0181-1916. doi: 10.1051/rnd:19840508.

14. Shuo Xiao, Jonathan R. Coppeta, Hunter B. Rogers, Brett C. Isenberg, Jie Zhu, Susan A. Olalekan, Kelly E. McKinnon, Danijela Dokic, Alexandra S. Rashedi, Daniel J. Haisenleder, Saurabh S. Malpani, Chanel A. Arnold-Murray, Kuanwei Chen, Mingyang Jiang, Lu Bai, Catherine T. Nguyen, Jiyang Zhang, Monica M. Laronda, Thomas J. Hope, Kruti P. Maniar, Mary Ellen Pavone, Michael J. Avram, Elizabeth C. Sefton, Spiro Getsios, Joanna E. Burdette, J. Julie Kim, Jeffrey T. Borenstein, and Teresa K. Woodruff. A microfluidic culture model of the human reproductive tract and 28-day menstrual cycle. Nature Communications, 8, mar 2017. ISSN 20411723. doi: 10.1038/ncomms14584.

15. J. E. Fléchon and R. H.F. Hunter. Distribution of spermatozoa in the utero-tubal junction and isthmus of pigs, and their relationship with the luminal epithelium after mating: A scanning electron microscope study. Tissue and Cell, 13(1):127–139, 1981. ISSN 00408166. doi: 10.1016/0040-8166(81)90043-4.

16. S S Suarez, I Revah, M Lo, and S Kölle. Bull sperm binding to oviductal epithelium is mediated by a Ca2+-dependent lectin on sperm that recognizes Lewis-a trisaccharide. Biology of reproduction, 59(1):39–44, jul 1998. ISSN 0006-3363. doi: 10.1095/biolreprod59.1.39.

17. Toru Hyakutake, Kotaro Mori, and Koichi Sato. Effects of surrounding fluid on motility of hyperactivated bovine sperm. Journal of Biomechanics, 71:183–189, apr 2018. ISSN 18732380. doi: 10.1016/j.jbiomech.2018.02.009.

18. Chih-kuan Tung, Lian Hu, Alyssa G. Fiore, Florencia Ardon, Dillon G. Hickman, Robert O. Gilbert, Susan S. Suarez, and Mingming Wu. Microgrooves and fluid flows provide preferential passageways for sperm over pathogen Tritrichomonas foetus. Proceedings of the National Academy of Sciences, 112(17):5431–5436, apr 2015. ISSN 0027-8424. doi: 10.1073/pnas.1500541112.

19. K. June Mullins and R. G. Saacke. Study of the functional anatomy of bovine cervical mucosa with special reference to mucus secretion and sperm transport. The Anatomical Record, 225(2):106–117, oct 1989. ISSN 10970185. doi: 10.1002/ar.1092250205.

20. Ting-Wei Su, Liang Xue, and Aydogan Ozcan. High-throughput lensfree 3D tracking of human sperms reveals rare statistics of helical trajectories. Proceedings of the National Academy of Sciences of the United States of America, 109(40):16018–22, oct 2012. ISSN 1091-6490. doi: 10.1073/pnas.1212506109.

21. Leonardo F. Urbano, Puneet Masson, Matthew VerMilyea, and Moshe Kam. Automatic Tracking and Motility Analysis of Human Sperm in Time-Lapse Images. IEEE Transactions on Medical Imaging, 36(3):792–801, mar 2017. ISSN 0278-0062. doi: 10.1109/TMI.2016.2630720.

22. Rupert P. Amann and Dagmar Waberski. Computer-assisted sperm analysis (CASA): Capabilities and potential developments. Theriogenology, 81(1):5–17, 2014. ISSN 0093691X. doi: 10.1016/j.theriogenology.2013.09.004.

23. P. Denissenko, V. Kantsler, D. J. Smith, and J. Kirkman-Brown. Human spermatozoa migration in microchannels reveals boundary-following navigation. Proceedings of the National Academy of Sciences, 109(21):8007–8010, may 2012. ISSN 0027-8424. doi: 10.1073/pnas.1202934109.

24. S. Fair, K. G. Meade, K. Reynaud, X. Druart, and S. P. De Graaf. The biological mechanisms regulating sperm selection by the ovine cervix, 2019. ISSN 17417899.

25. Luis Alvarez, Benjamin M. Friedrich, Gerhard Gompper, and U. Benjamin Kaupp. The computational sperm cell. Trends in Cell Biology, 24(3):198–207, 2014. ISSN 09628924. doi: 10.1016/j.tcb.2013.10.004.

26. Susan S. Suarez. Mammalian sperm interactions with the female reproductive tract. Cell and Tissue Research, 363(1):185–194, jan 2016. ISSN 14320878. doi: 10.1007/s00441-015-2244-2.

27. Laura M King, Denise R Holsberger, and Ann M Donoghue. Correlation of CASA Velocity and Linearity Parameters With Sperm Mobility Phenotype in Turkeys. Journal of Andrology J Androl, 2121(1):65–71, jan 2000. ISSN 0196-3635. doi: 10.1002/j.1939-4640.2000.tb03277.x.

28. P E Kloeden and E Platen. Numerical Solution of Stochastic Differential Equations, volume 23. Springer Berlin Heidelberg, Berlin, Heidelberg, 1995. ISBN 978-3-642-08107-1. doi: 10.1007/978-3-662-12616-5.

29. H. J. Schuberth, U. Taylor, H. Zerbe, D. Waberski, R. Hunter, and D. Rath. Immunological responses to semen in the female genital tract. Theriogenology, 70(8):1174–1181, 2008. ISSN 0093691X. doi: 10.1016/j.theriogenology.2008.07.020.

30. Kiyoshi Miki and David E. Clapham. Rheotaxis guides mammalian sperm. Current Biology, 23(6):443–452, mar 2013. ISSN 09609822. doi: 10.1016/j.cub.2013.02.007.

31. M. Burkitt, D. Walker, D. M. Romano, and A. Fazeli. Using computational modeling to investigate sperm navigation and behavior in the female reproductive tract. Theriogenology, 77 (4):703–716, mar 2012. ISSN 0093691X. doi: 10.1016/j.theriogenology.2011.11.011.

32. Justus A. Kromer, Steffen Märcker, Steffen Lange, Christel Baier, and Benjamin M. Friedrich. Decision making improves sperm chemotaxis in the presence of noise. PLOS Computational Biology, 14(4):e1006109, apr 2018. ISSN 1553-7358. doi: 10.1371/journal.pcbi.1006109.

33. W Schroeder, K Martin, and B Lorensen. The visualization toolkit: an object oriented approach to 3D graphics. New York: Kitware. Kitware, 2006. ISBN 9781930934191.

34. Utkarsh Ayachit. The ParaView Guide - Updated for ParaView version 4.3. 2015. ISBN 9781930934306 1930934300.

35. Byung Chan Eu. Generalization of the Hagen–Poiseuille velocity profile to non-Newtonian fluids and measurement of their viscosity. American Journal of Physics, 58(1):83–84, jan 1990. ISSN 0002-9505. doi: 10.1119/1.16328.

36. Busch and Waberski. Künstliche Besamung bei Hausund Nutztieren. Schattauer, 2007. ISBN 3-334-00379-5.

37. J M Cummins and P F Woodall. On mammalian sperm dimensions. Journal of reproduction and fertility, 75(1):153–175, sep 1985. ISSN 1470-1626. doi: 10.1530/jrf.0.0750153.

